# Differential Roles of Positive and Negative Valence Systems in the Development of Post-Traumatic Stress Psychopathology

**DOI:** 10.1101/2021.03.08.434335

**Authors:** Ziv Ben-Zion, Ofir Shany, Roee Admon, Nimrod Jackob Keynan, Netanell Avisdris, Shira Reznik Balter, Arieh Y. Shalev, Israel Liberzon, Talma Hendler

## Abstract

Negative and positive valence systems (NVS and PVS) pertain to processing of aversive and rewarding stimuli, respectively. Post-Traumatic Stress Disorder (PTSD) has been typically associated with hyper-responsivity of the NVS, and more recently, with deficient PVS functionality. The respective roles of these systems in early PTSD development have yet to be resolved. Here, we assessed neurobehavioral indicators of PVS and NVS longitudinally among 171 adult civilians at 1-, 6-, and 14-months following trauma exposure (TP1, TP2, and TP3). Using the ‘Safe or Risky Domino Choice’ (SRDC) game during fMRI, NVS and PVS functionality (i.e., activity and connectivity) were indicated by the amygdala and ventral striatum (VS) responses to punishments and rewards, respectively. The complementary functionality of these systems was behaviorally assessed as the percentage of risky choices taken during the game. Results revealed that increased amygdala functionality at TP1 was associated with greater PTSD severity at TP1 and TP3, specifically with hyperarousal and intrusion. Decreased VS functionality at TP1 was associated with greater PTSD severity at TP3, specifically with avoidance. Explainable machine learning revealed the primacy of PVS over NVS functionality at TP1 in predicting PTSD severity at TP3. Behaviorally, fewer risky choices were associated with more severe symptoms at TP1, especially with intrusion and avoidance. Overall, these results suggest a differential and potentially complementary involvement of NVS and PVS in PTSD development following trauma. Early therapeutics for PTSD in the immediate aftermath of trauma may thus target both negative and positive valence processing.

## Introduction

The concept of separate systems for the processing of negative and positive valence stimuli (NVS and PVS, respectively) originated in psychology over a century ago, yet more recently was incorporated into the field of clinical neuroscience^1^. These systems were further identified as two core dimensions of human behavior in the NIMH Research Domain Criteria (RDoC)^2,3^. The NVS mediates responses to aversive situations or contexts, evoking negative feelings such as fear, anxiety, and loss, whereas the PVS mediates responses to positive motivational situations or contexts such as reward-seeking, consummatory behavior, and reward learning. Valence estimation could be challenging in real-life as stimuli often evoke mixed or even conflicting emotions and consequence behaviors. Further challenge for these systems may stem from the exposure to traumatic stress. Indeed, stress was shown to hinder accurate valence estimations^4–6^, as it increases vigilance and drains cognitive resources^7,8^. While such restrictions at the immediate aftermath of traumatic events might be beneficial for survival, a transition into reward-driven behavior over time, despite the presence of a heightened threat, is necessary for promoting stress resilience^9–13^. Chronic stress psychopathologies such as Post-Traumatic Stress Disorder (PTSD) are often characterized by a tendency to sacrifice potential rewards in order to avoid aversive encounters^14–17^. The respective roles of NVS and PVS in the early development of PTSD has yet to be investigated. Here, we examined the idea that individuals’ ability to recover from a traumatic event relies on differential processing of the PVS and NVS, by assessing neurobehavioral manifestations of these systems separately and together, in the early aftermath of trauma.

Substantial evidence links PTSD to abnormal NVS functionality, consistently showing increased sensitivity to various negative stimuli (e.g. symptom provocation, fearful faces)^18,19^ among PTSD patients, potentially reflecting clinical symptoms of hyperarousal and intrusion (i.e., re-experiencing)^20–23^. Furthermore, studies using decision-making tasks demonstrated an association between PTSD and increased behavioral aversion to risk^24,25^ and ambiguous losses^26^, possibly reflecting the generalized (not only trauma-specific) oversensitivity of the NVS, often seen in PTSD patients. At the neural level, the role of the NVS in PTSD has been repeatedly documented as abnormally heightened amygdala activation and hyperactive salience network (e.g., anterior insula, dorsal anterior cingulate cortex) in response to negative stimuli^20,27–29^. Furthermore, aberrant functional connectivity of the amygdala with the prefrontal cortex (PFC) in response to negative stimuli was observed in PTSD, specifically with the orbitofrontal cortex (OFC)^30,31^, suggesting disrupted emotion regulatory capacity.

More recent work suggests that PTSD might also involve abnormalities in the PVS, as indicated by deficient reward anticipation, decreased approach (reward-seeking) behavior, and diminished hedonic responses to rewarding outcomes^32,33^. Reward processing is known to involve the meso-corticolimbic pathway, represented by dopamine projections from the ventral tegmental area (VTA) to the ventral striatum (VS), including the nucleus accumbens (NAcc), and further to ventromedial/orbital frontal brain structures^34,35^. While decreased VS activation to positive stimuli was demonstrated initially in depressed individuals mostly in relation to anhedonia^36,37^, it was also reported in PTSD patients in response to monetary gains^38,39^ and happy faces^40^. Recent studies further pointed to aberrant functional connectivity between the VS and the ventromedial PFC (vmPFC) in PTSD, suggesting altered function of the reward circuitry in this disorder^41,42^.

Although both systems seem to have a role in PTSD, their relative contribution to the development of post-traumatic psychopathology remains largely unknown, due to several substantial clinical and methodological challenges. First, only a small portion (around 20%) of those with early stress symptoms go on to develop chronic PTSD^43,44^. Second, even within the group of PTSD patients, clinical phenotypes are heterogeneous^45,46^, and different symptom manifestations (e.g. hyperarousal vs. avoidance) might be related to different neurobehavioral processes (e.g. NVS vs. PVS). Third, the typical cross-sectional designs used in PTSD research cannot infer on the immediate response to trauma nor on any potential dynamics that may occur during the first year post-trauma, a critical period that determines who will develop PTSD and who will recover^47,48^. While recent years show an increase in longitudinal studies, the majority focused solely on either NVS or PVS, thus cannot infer on the unique role of each system in PTSD development over time.

To overcome these challenges, we conducted a large-scale longitudinal fMRI study of recent trauma survivors (see study protocol^49^). A sample of n=171 adult civilians were screened for early PTSD symptoms, suggestive of chronic PTSD risk^50,51^, within 10-14 days following their release from a general hospital’s emergency room (ER). Participants were longitudinally tracked at 1-, 6- and 14-months following trauma exposure (TP1, TP2, and TP3, respectively) as they underwent fMRI scanning while playing ‘Safe of Risky Domino Choice’ (SRDC) game. The SRDC is a naturalistic interactive gambling game encompassing safe or risky choices resulting in outcomes of reward (win) or punishment (loss). Based on previous studies using the SRDC game^25,52–55^, NVS and PVS functionality (i.e., activity and connectivity) were indicated by the amygdala and ventral striatum (VS) responses to punishments and rewards, respectively. In order to win in the SRDC game, individuals’ behavior must include both safe and risky choices, enabling a behavioral marker of the complementary functionality of the PVS and NVS.

**Our first aim** was to establish the link between neural indicators of NVS and PVS and post-traumatic symptom severity shortly after exposure (TP1). Based on previous findings^25,56^, we hypothesized that more severe post-traumatic symptoms would be associated with greater response of the amygdala to punishments, decreased response of the VS to rewards, and altered functional connectivity of the VS and the amygdala with the PFC. **Our second aim** was to reveal the predictive contribution of early NVS and PVS neural functionality to PTSD symptom development within the first year following trauma exposure. We hypothesized that greater activity and connectivity of the amygdala to punishments and decreased activity and connectivity of the VS to rewards at TP1 would be predictive of more severe post-traumatic symptoms at TP2 and TP3 (beyond initial severity at TP1). By utilizing an explainable machine learning, we further examined the relative importance of PVS and NVS neural processing at TP1 to PTSD severity at TP3. **Our third aim** was to unveil the complementary role of NVS and PVS in PTSD symptom development through behavioral indices in the SRDC game. Based on previous work^25^, we hypothesized that decreased risk-taking behavior at TP1 would be associated with more severe symptoms at TP1, TP2 and TP3, possibly reflecting increased threat from punishments (i.e., hyperactive NVS) and decreased hedonic response to rewards (i.e., hypoactive PVS).

## Methods

### Participants

The study group included adult survivors of traumatic events who were admitted to Tel-Aviv Sourasky Medical Center’s Emergency Room (ER). The most common trauma type among participants was motor vehicle accidents (n=137, 80%), while other traumatic events included assaults, terror attacks, drowning, mass casualty incidents, animal attacks, robbery, and electrocution. Exclusion criteria included head trauma or coma exceeding 30 minutes, incompatibility for MRI scanning, history of substance abuse, current or past psychotic disorder, or chronic PTSD diagnosis pre-admission to ER. Only trauma survivors without a known medical condition that interfered with their ability to give informed consent or to cooperate with screening and/or treatment were included. A total of 171 participants completed clinical and neuroimaging assessments within one-month following their traumatic incident (TP1). Of these, 39 individuals were excluded due to missing (n=16) or partial (n=5) functional scan while performing the SRDC game; poor quality of the functional scan (e.g., movements, artifacts, etc.) (n=6); missing or poor structural scan (n=5); missing or partial behavioral data of the paradigm (n=5); not understanding the instructions properly (n=1); and missing clinical data (n=1), for a final sample size of 132 individuals at TP1. Of these 132 participants, 115 and 112 participants completed clinical interviews at TP2 and TP3 (respectively). Participants’ demographic and clinical characteristics across the three time-points (TP1, TP2, and TP3) are presented in Table 1. Of note, 6 individuals were excluded from the neural analysis at TP2 and 22 at TP3, due to missing or partial or poor quality of the functional scan, missing / poor quality structural scan, or missing/partial behavioral data of the SRDC game, for a final sample size of 109 and 90 individuals with valid neural data at TP2 and TP3, respectively.

**Table 1.** Participants’ demographic and clinical characteristics. The table summarizes characteristics of participants included in the final analyses across the three time-points. Means and standard deviations of participants’ age, gender (Female: Male), and PTSD severity (CAPS-5 total scores), at 1-, 6- and 14-months post-trauma (TP1, TP2 and TP3). Additionally, the percentage of motor-vehicle accidents of individuals diagnosed with PTSD (%MVA’s, %PTSD) are reported for each time-point separately.

| **Measure** | <u>TP1 (n=132)</u> |  | <u>TP2 (n=115)</u> |  | <u>TP3 (n=112)</u> |  |
| --- | --- | --- | --- | --- | --- | --- |
|  | *M* | *SD* | *M* | *SD* | *M* | *SD* |
| Age | 33.52 | 11.01 | 33.73 | 11.08 | 33.56 | 11.27 |
| Gender (F: M) | 63: 69 | - | 55: 60 | - | 56: 56 | - |
| CAPS-5 Total | 24.91 | 11.68 | 14.97 | 10.89 | 10.69 | 10.10 |
| % MVA's | 89% (n=117) |  | 88% (n=101) |  | 88% (n=99) |  |
| % PTSD | 74% (n=97) |  | 35% (n=40) |  | 24% (n=27) |  |

### Procedure

A member of the research team identified potential trauma-exposed individuals via the ER computerized medical records. Within 10–14 days of trauma exposure, approximately n=4,000 potential participants were contacted by telephone for initial screening. Acute PTSD symptoms (indicative of the risk for PTSD development^50^) were assessed using a modified dichotomous version of the PTSD Checklist (PCL) questionnaire^57^. Those who met PTSD symptom criteria (except for the “1-month duration” criteria) and did not meet any of the exclusion criteria (see under Participants), were invited to participate to a face-to-face clinical assessment and an fMRI scan, at one-month post-trauma (TP1). We preferentially enrolled survivors who met PTSD diagnosis in the face-to-face clinical interview, but also enrolled some individuals with sub-threshold PTSD (n=35). Two identical follow-up meetings, including both clinical and neural assessments, were conducted at 6-and 14-months following trauma (TP2 and TP3, respectively). For more details, see Ben Zion et al. (2019).^49^

### Clinical Assessments

PTSD diagnosis and severity at each time-point were determined by a comprehensive clinical interview conducted by trained and certified clinical interviewers. A continuous measure of total symptom severity was obtained by summing individual items’ scores of the Clinician-Administered PTSD Scale for DSM-5 (CAPS-5)^58^, the current gold-standard for PTSD diagnosis. Total scores were further computed for each of the DSM-5 symptom clusters: intrusion (cluster B), avoidance (cluster C), negative alterations in cognition and mood (cluster D), and hyperarousal (cluster E).

### fMRI Paradigm: Safe or Risky Domino Choice (SRDC) game

During scanning, participants played a 2-player gambling game for 14 minutes, in which they were required to take risky choices in order to win (see Fig. 1). The effectiveness of the SRDC to detect individuals’ sensitivity to risk, punishment and reward was previously validated in both healthy and clinical populations^25,52–55,59^. Participants were told that their opponent is the experimenter who decides whether to expose their choice or not, thus their choices can increase their chances of winning. In fact, the computer randomly generated the opponent’s responses in a predetermined pattern to allow a balanced design (exposing the player’s choices 50% of the time). This paradigm involve four intervals (Fig. 1): <u>decision-making</u> (deciding which chip to choose), <u>decision-execution</u> (taking risky or safe chips), <u>anticipation of an outcome</u> (waiting passively for the opponent’s decision to expose or not to expose player’s choice), and <u>response to an outcome</u> (receiving a reward or punishment, defined by the chosen chip and opponent’s decision). Here, we focused on the decision-making interval for behavioral indexing (i.e., individual tendency to take risky vs safe choices) and on the neural responses in the anticipation for an outcome interval (i.e., responses to rewards vs. punishments). At the beginning of each game, 12 random domino chips were assigned to the participant, while one domino “master” chip (constant throughout the game) appeared at the top left corner of the board. In each round of the game, players had to choose one chip and place it face down adjacent to the master chip. They then had to wait for the opponent’s response (i.e., anticipation), to see whether the opponent challenges this choice by uncovering the chosen chip or not (i.e., outcome). Players were able to win this competitive game if they successfully disposed all 12 chips within 4 minutes. Each assigned chip either matched the master chip (had at least one of the master chip’s numbers) or did not match. Since the master chip was constant throughout the game, it was only possible to win by choosing both matching and non-matching chips. In the game context, matching chips were considered ‘‘safe’’ moves since they were associated with rewards, while non-matching chips were considered ‘‘risky’’ moves since they were associated with punishments. Accordingly, based on players’ choices, there were two possible anticipation periods, “risky anticipation” following a non-matching choice or “safe anticipation” following a matching choice. Based on players’ choices and opponent’s response, there were four possible consequences (outcomes) per game round: (1) <u>Show of a non-matching chip (i.e., main punishment):</u> the choice of a non-matching chip was exposed and the player was punished by receiving back the selected chip plus 2 additional chips from the deck; <u>(2) No-show of a non-matching chip (i.e., relative reward):</u> the choice of a non-matching chip remained unexposed and only the selected chip was disposed of, so the player was relatively rewarded as they got away with a non-matching choice; <u>(3) Show of a matching chip (i.e., main rewards):</u> the choice of a matching chip was exposed and the player was rewarded by the disposal of the selected chip and one additional random chip from the game board; <u>(4) No-show of a matching chip (i.e., relative punishment):</u> the choice of a matching chip was not exposed and only the selected match chip was disposed of, so the player was relatively punished as they could have disposed of a non-matching chip instead. For more details, see supplementary methods.

**Figure 1.**
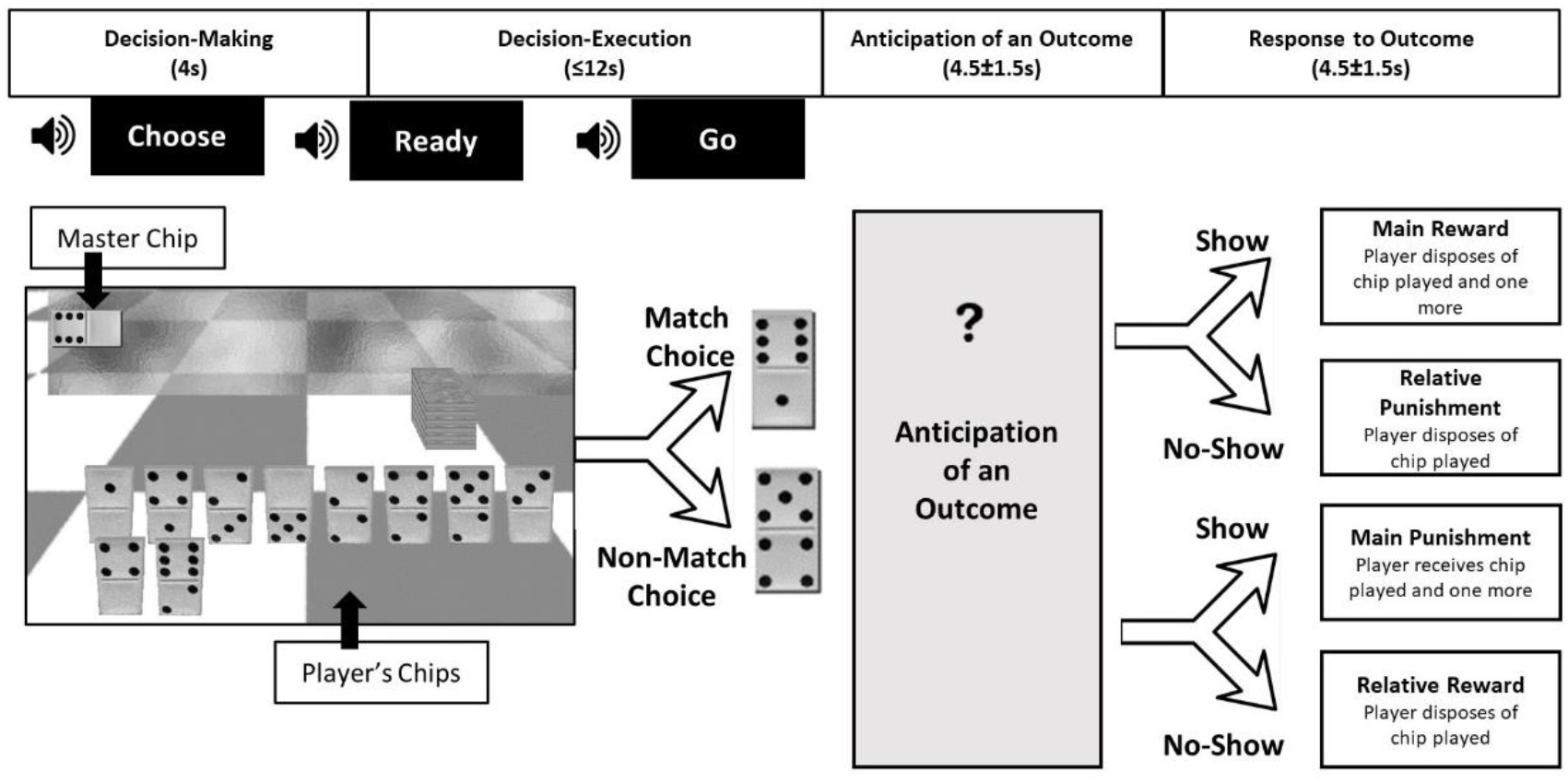
Safe or Risky Domino Choice (SRDC) Paradigm. Each round of the game is composed of four intervals. First, participants choose which chip to play next (i.e., decision-making), either a matching choice (e.g., a chip with least one of the master chip’s numbers) or a non-matching choice. Next, they move the cursor to the chosen chip and place it facing down adjacent to the master chip (i.e., decision-execution). Participants then wait for the opponent’s response (i.e., anticipation of an outcome) to see whether the opponent challenges their choice by uncovering the chosen chip or not (i.e., response to outcome). Participants’ choices and opponents’ responses are interactively determined by the flow of the game round after round, creating a natural progression of a game situation that lasts 4 min or until the player wins the game by disposing of all his chips. Each player played consecutively for 14 min (approximately 3-4 game rounds).

### Behavioral Analysis of SRDC game

To characterize individuals’ behavioral choices during the game, a ‘risky choice index’ was defined as the ratio between the number of risky choices (e.g., choosing a non-matching chip) and the total number of choices throughout the entire game (e.g., choosing either a matching or non-matching chip), multiplied by 100 (to obtain percentage). Rounds in which participants had no actual choice between safe and risky choices were excluded (i.e., when there were only matching or only non-matching chips on the screen). This index represents a nonbiased choice when equal to 50% (exactly half of the choices were non-matching chips), a bias towards riskier behavior when greater than 50%, and a bias towards safer behavior (i.e., risk aversion and avoidance tendencies) when less than 50%.

### fMRI Data Acquisition

Whole-brain functional and anatomical images were acquired using a 3.0 Tesla Siemens MRI system (MAGNETOM Prisma, Germany) with a 20-channel head coil at Tel-Aviv Sourasky Medical Center. Functional images were acquired in an interleaved order (anterior to posterior), using a T2*-weighted gradient-echo planar imaging pulse sequence (TR/TE=2000/28ms, flip angle= 90°, voxel size 2.2mm^3^, FOV=220×220mm, slice thickness=3mm, 36 slices per volume). A T1-weighted three-dimensional anatomical image was also collected, using a magnetization prepared rapid gradient echo (MPRAGE) sequence (TR/TE=2400/2.29ms, flip angle = 8°, voxel size 0.7 × 0.7 × 0.7 mm, FOV = 224 × 224 mm), enabling optimal localization of the functional effects.

### fMRI Data Analysis

Preprocessing was conducted using FMRIPREP version 1.5.8^60^, a Nipype based tool^61^. The preprocessing procedures included spatial realignment of the echo planar imaging volumes, co-registration with the structural image, segmentation, normalization, and spatial smoothing (for full details, see supplementary methods). Preprocessed imaging data was analyzed using Statistical Parametric Mapping 12 (SPM12). The size of the effect for each condition for each participant was computed by a general linear model (GLM) that included the different conditions of the game: “choose”, “ready”, “go”, “picked match”, “picked non-match”, “show match”, “show non-match”, “no show match” and “no show non-match”. Individual statistical parametric maps were calculated for the a-priori defined contrasts of interest of “rewards vs. punishments” (receiving both rewarding outcomes > receiving both punishing outcomes) and “punishments vs. rewards” (receiving both punishing outcomes > receiving both rewarding outcomes). This was done for all participants at each time-point separately (n=132 at TP1, n=109 at TP2, and n=90 at TP3).

#### Whole-Brain and ROI analysis

Based on previous findings using the same paradigm^25,52–55,59^, two main regions of interest (ROIs) were defined using the Human Brainnetome atlas (Fan and colleagues^62^) and California Institute of Technology 168 (CIT-168) atlas (Pauli and colleagues^63^). In the contrast of rewards vs. punishments, the focus was on the left and right ventral striatum (VS), composed of the ventral caudate from the Human Brainnetome atlas (regions 219-220) and the nucleus accumbens from the CIT-168 atlas. In the opposite contrast of punishments vs. rewards, the focus was on the left and right amygdala, composed of the medial (regions 211-212) and lateral amygdala (regions 213-214) from the Human Brainnetome atlas. MarsBaR region of interest toolbox for SPM^64^ was used to extract participants’ contrast activations (average beta weight) from each ROI separately for each time point. Analyses were performed separately for each hemisphere (left and right amygdala, left and right VS). For more details regarding whole-brain and ROI analyses, see supplementary methods.

#### Functional Connectivity Analysis

Examination of functional connectivity interactions with the task regressors was performed using generalized psychophysiological interaction (gPPI) as implemented in CONN toolbox^65,66^ (for full details, see supplementary methods). ROI to ROI analysis was performed using the two main a-priori ROIs as seed regions -amygdala and VS -and two a-priori selected regions within the prefrontal cortex as target regions -vmPFC and lateral OFC (lOFC). The selection of these two regions was based on extensive literature pointing to involvement of the human PFC and OFC in processing reward and punishment. The two target regions were anatomically defined by the Human Brainnetome atlas^62^: The vmPFC was composed of regions 41-42, 47-48, whereas the lOFC was composed of regions 43-44, 45-46, 51-52. For more details, see supplementary methods.

### Statistical Analysis

IBM SPSS Statistics for Windows^67^ and R^68^ software were both used for the statistical procedures. Participants with extreme scores of ±3 standard deviations from the mean in the relevant variables were defined as outliers and excluded from the analysis. For all statistical tests, α=0.05 was used with either one-sided a-priori hypotheses or two-sided non-directional hypotheses (according to the a-priori hypotheses outlined at the end of the introduction section). Benjamini–Hochberg False Discovery Rate (FDR) correction (q<0.05)^69^ was calculated to control for multiple comparisons.

### Predictor Importance Ranking

To examine the contribution of early neural activations (at TP1) and rank their importance for the prediction of PTSD symptom severity at the study’s endpoint (TP3), we used SHAP (Shapley Additive exPlanation)^70^, a state-of-the-art methodology in the field of explainable machine learning. SHAP estimates “Shapely” values, which provide a surrogate for the individual additive contribution of each feature to the prediction. In other words, SHAP’s rank order informs which feature values mostly influence the prediction, while accounting for the influence of all other feature values, and while controlling for the order in which features are added to the model^70^.

## Results

### Neural Indicators of PVS and NVS Associated with PTSD Symptom Severity Shortly after Trauma Exposure (TP1)

Partial correlations were computed between neural indices (mean activations of the VS and amygdala to rewards vs. punishments, as well as their connectivity patterns), and CAPS-5 total scores at TP1, while controlling for participants’ age, gender, and trauma type (i.e., covariates). Results revealed a significant positive correlation between response to punishments (vs. rewards) in the amygdala and symptom severity at TP1 (n=128; left amygdala: r=0.155, p=0.043, pFDR=0.043; right amygdala: r=0.162, p=0.035, p_FDR_=0.043). Even further, examining functional connectivity patterns of the amygdala with predetermined PFC regions (vmPFC and lOFC) revealed that stronger amygdala-lOFC connectivity in response to punishments (vs. rewards) was associated with more severe PTSD symptoms (n=124; right amygdala – left lOFC: r=0.254, p=0.005, q_-FDR_<0.05). Taken together, severity of post-traumatic symptoms shortly after exposure was positively associated with amygdala activation and amygdala-lOFC connectivity in response to punishment (Fig. 2 a&c). Contrary to our expectation, VS activation in response to rewards (vs. punishments) was not significantly associated with symptom severity at TP1 (n=131; left VS: r=0.022, p=0.401; right VS: r=0.048, p=0.297; Fig. 2b), nor does VS functional connectivity with the predetermined PFC regions in response to rewards (vs. punishments) (n=122, q_-FDR_>0.05). For whole-brain results, please refer to supplementary results, supplementary table 1, and supplementary figure 1.

**Figure 2.**
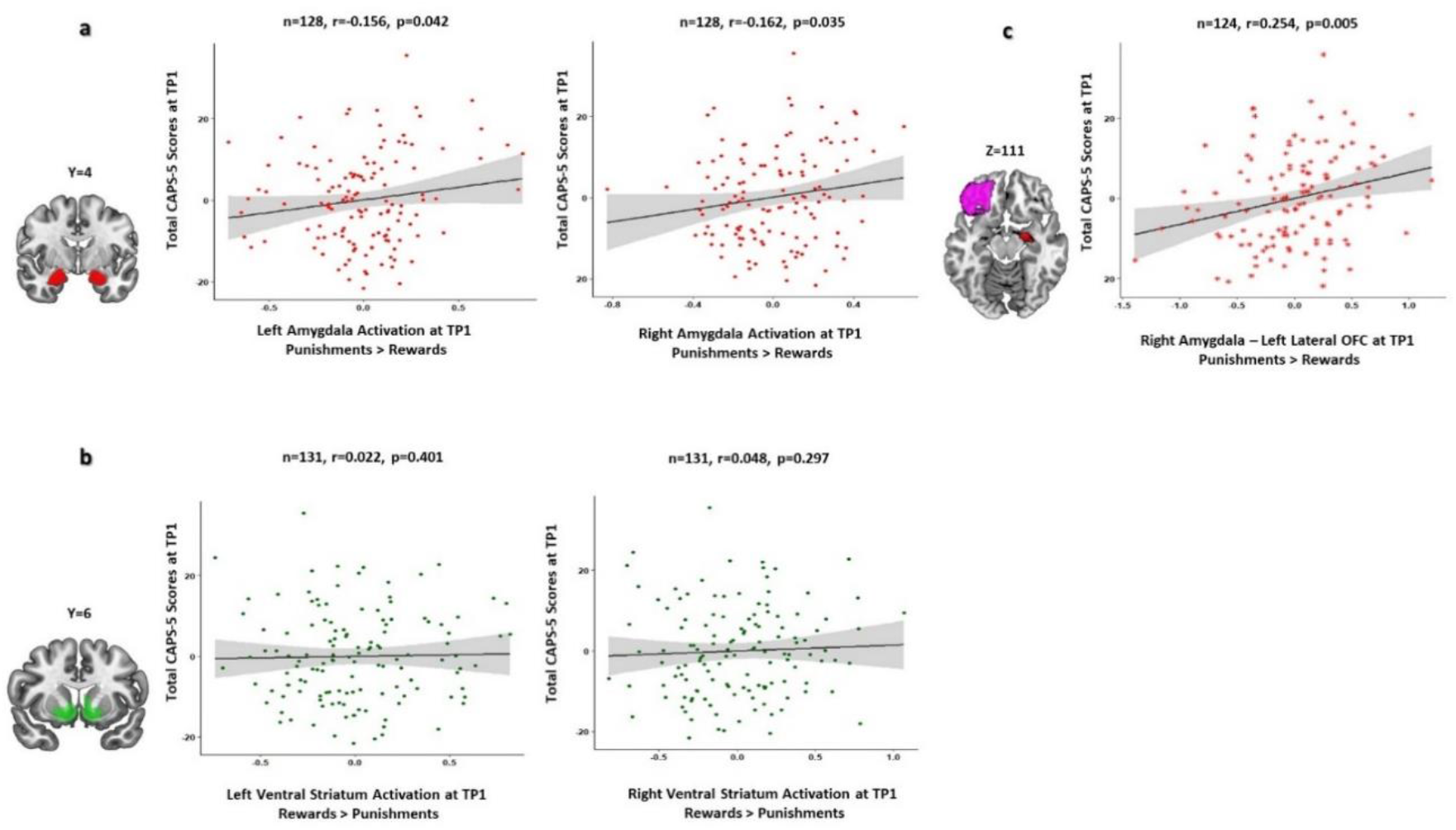
Neural Indicators of PVS and NVS Associated with PTSD Symptom Severity Shortly after Trauma Exposure (TP1). **a**. Partial regression scatter plots depicting the relation between CAPS-5 total scores at TP1 (y-axis) and activation (mean beta values) of the left and right amygdala in response to punishments vs. rewards (x-axis). The anatomical amygdala ROI which was used for this analysis is presented on a coronal view of the brain (in red). Each dot represents one subject. **b**. Partial regression scatter plots depicting the relation between CAPS-5 total scores at TP1 (y-axis) and activation (mean beta values) of the left and right ventral striatum in response to rewards vs. punishments (a-axis). The anatomical ventral striatum ROI which was used for this analysis is presented on a coronal view of the brain (in green). Each dot represents one subject. **c**. Partial regression scatter plots depicting the relation between CAPS-5 total scores at TP1 (y-axis) and connectivity (mean beta values) of the right amygdala – left lateral OFC in response to punishments vs. rewards at TP1 (x-axis). The anatomical ROIs which were used for this analysis, right amygdala (red) and left lateral OFC (violet), are presented on an axial view of the brain. Each asterisk represents one subject. Values on all axes are unstandardized residuals.

### Neural Indicators of PVS and NVS Shortly after Exposure (TP1) Predict PTSD Symptom Severity One Year Following Trauma (TP3)

To assess the relationship between NVS and PVS functionality shortly after exposure and PTSD severity within the first year following trauma exposure, partial correlations were computed between neural indices at TP1 and CAPS-5 total scores at TP2 and TP3, while controlling for participants’ age, gender, trauma type, and initial symptom severity (CAPS-5 total scores at TP1). In line with our hypothesis, both hyperactive NVS (i.e., amygdala’s activation to punishment) and hypoactive PVS (i.e., VS activation to reward) at TP1 were significantly predictive of more severe PTSD symptoms at TP3. Specifically, greater left amygdala activation at TP1 was associated with higher CAPS-5 total scores at TP3 (n=108; r=0.197, p=0.022) (see Fig. 3a); and decreased right VS activation at TP1 was associated with higher CAPS-5 total scores at TP3 (n=111, r=-0.235, p=0.007) (see Fig. 3b). Individuals with hyperactive NVS or hypoactive PVS early after trauma (TP1) were thus prone to develop more severe symptoms a year later (TP3), beyond their initial symptom severity (at TP1). Testing specific symptom clusters, greater amygdala activation to punishments at TP1 was associated with more severe hyperarousal (r=0.176, p=0.037, p_FDR_=0.074) and intrusion at TP3 (r=0.217, p=0.027, p_FDR_=0.074; Fig. 3a), whereas decreased VS activation to rewards at TP1 was linked to more avoidance at TP3 (r=-0.285, p=0.001, p_FDR_=0.004; Fig. 3b).

**Figure 3.**
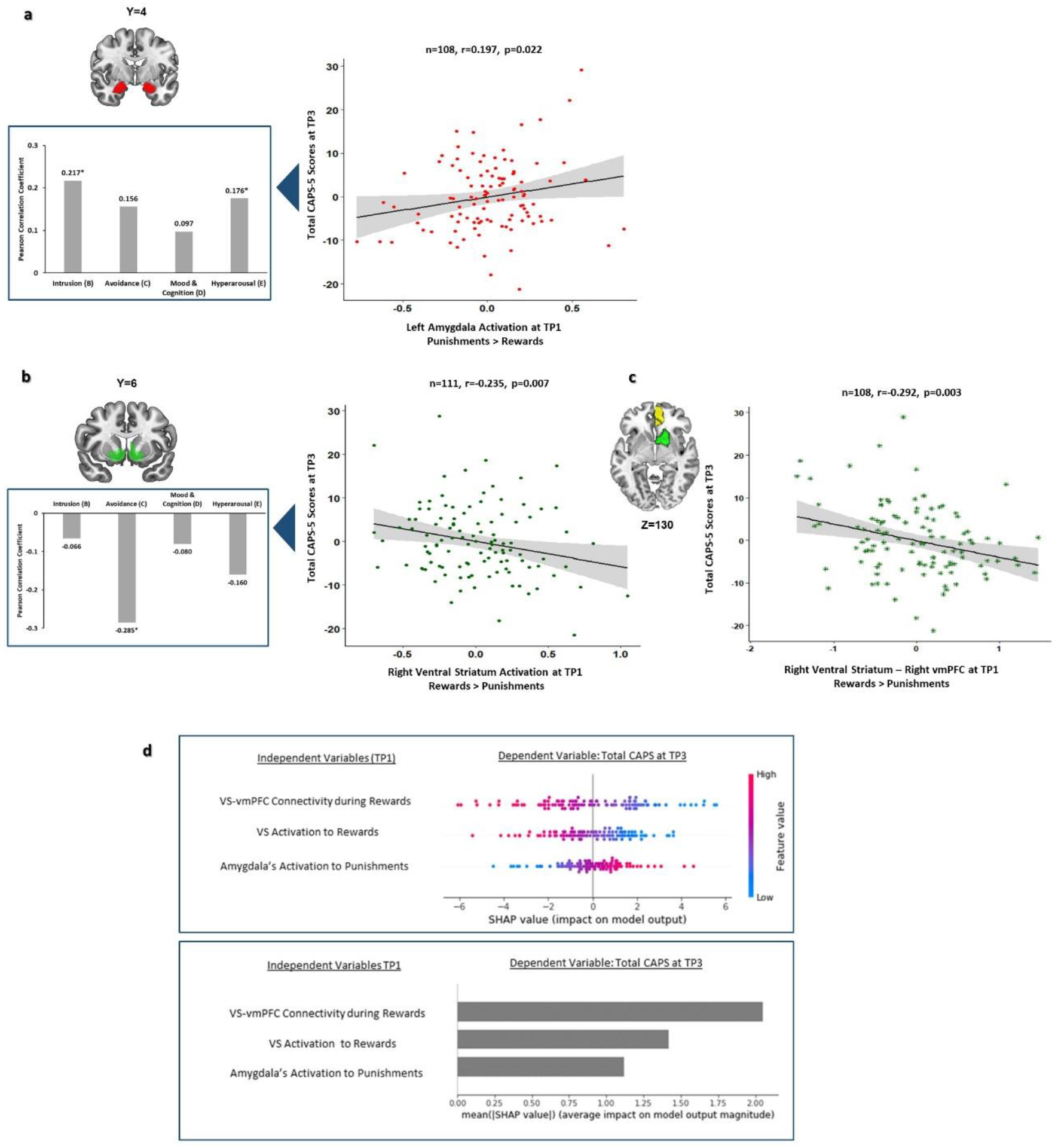
Neural Indicators of PVS and NVS Shortly after Exposure (TP1) Predict PTSD Symptom Severity One Year Following Trauma (TP3). **a**. Partial regression scatter plot depicting the relation between CAPS-5 total scores at TP3 (y-axis) and activation (mean beta values) of the left amygdala in response to punishments vs. rewards at TP1 (x-axis), while controlling for age, gender and trauma type (covariates). Values on both axes are unstandardized residuals. On the left, a bar plot presenting the correlations between left amygdala activation and all four PTSD symptom clusters at TP3 according to CAPS-5: intrusion (B), avoidance (C), negative alterations in cognition and mood (D), and hyperarousal symptoms (E). Pearson correlation coefficients (r) are presented above each bar. *p-FDR<0.05. **b**. Partial regression scatter plot depicting the relation between CAPS-5 total scores at TP3 (y-axis) and activation (mean beta values) of the right ventral striatum (VS) in response to rewards vs. punishments at TP1 (x-axis), while controlling for covariates (see in a). On the left, a bar plot presenting the correlations between right VS activation and all four PTSD symptom clusters at TP1 according to CAPS-5. Pearson correlation coefficients (r) are presented above each bar. *p-FDR<0.05. **c**. Partial regression scatter plot depicting the relation between CAPS-5 total scores at TP3 (y-axis) and connectivity (mean beta values) of the right VS – right vmPFC in response to rewards vs. punishments at TP1 (x-axis), while controlling for covariates (see in a). The corresponding predefined anatomical ROIs, right VS (green) and right vmPFC (yellow), are presented next to the plot.**d. Explainable machine learning**. Top panel - absolute feature importance as calculated by SHAP, pointing to the importance of neural features at TP1 in predicting CAPS-5 total scores at TP3. The larger the SHAP value, the more important the feature is to discriminate between individuals with different symptom severity (CAPS-5 total scores). Bottom panel SHAP importance summary dot plot displaying features that influenced the linear regression model predictions of PTSD symptom severity (CAPS-5 total scores) at TP3. Features are first sorted by their global impact (y-axis). For every individual from the n=105 included in our sample, a dot represents the attribution value for each feature from low (blue) to high (red).

Of note, neither amygdala activation to punishments nor VS activation to rewards at TP1 were significantly associated with CAPS-5 total scores at TP2 (n=114; left amygdala: r=-0.021, p=0.413; right amygdala: r=-0.146, p=0.320; left VS: r=0.065, p=0.249; right VS: r=0.006, p=0.475), nor with any specific CAPS-5 symptom clusters at TP2.

Examining the predictive power of functional connectivity patterns of the neural components of the two valence systems at TP1 for predicting symptom severity at TP3 revealed such a relation only for neural components of the PVS. Specifically, decreased VS-vmPFC connectivity during rewards (vs. punishments) at TP1 was associated with more severe post-traumatic stress symptoms at TP3 (n=108, right VS – right vmPFC: r=-0.292, p=0.003, q_-FDR_<0.05), indicating that individuals with less VS-vmPFC connectivity at TP1 developed more severe PTSD symptoms at TP3 (Fig. 3c). Amygdala functional connectivity with the predetermined PFC regions (vmPFC and lOFC) in response to punishments (vs. rewards) at TP1 was not related to PTSD symptom severity at TP3 (n=110, q_-FDR_>0.05).

Finally, we tested the relative contribution of amygdala and VS functionality (activation and connectivity) at TP1 for PTSD symptom severity per CAPS-5 scores at TP3, using linear regression with the baseline neurobehavioral indices of PVS and NVS from TP1, which significantly predicted post-traumatic stress symptoms at TP3: left amygdala activation to punishments (vs. rewards) (Fig. 3a), right VS activation to rewards (vs. punishments) (Fig. 3b), and right VS–right vmPFC functional connectivity during rewards (vs. punishments) (Fig. 3c). As expected, all three variables together at TP1 accounted for a significant amount of variance of CAPS-5 total scores at TP3 (n=105, R^2^=0.200, F_3,101_=8.398, p<0.001). To identify the relative importance of these three neural predictors, we calculated importance values using the SHAP^70^ analytic approach (see Predictor Importance Ranking in Methods). In terms of absolute feature importance, PVS abnormalities showed greater importance compared to NVS abnormalities. To this end, VS-vmPFC connectivity during rewards (vs. punishments) at TP1 was the best predictor of subsequent CAPS scores at TP3, followed by the right VS response to rewards (vs. punishments) at TP1, and lastly left amygdala response to punishments (vs. rewards) at TP1 (Fig. 3d). Notably, when comparing the variances of VS activity and connectivity to amygdala activity, it seems that while the importance/contribution of the VS differed greatly between individuals (SHAP values ranging from -6 to +6), the amygdala had a small contribution in most participants (SHAP values between -2 to +2), and a large contribution to only a minority (Fig. 3d).

### Behavioral Indicators for the Complimentary Role of PVS and NVS in PTSD Symptom Development

Partial correlations were computed between risky choice index at TP1 and CAPS-5 total scores at TP1, TP2, and TP3, while controlling for participants’ age, gender, and initial symptom severity. In line with our hypothesis, a significant negative correlation was found between risky choice index and CAPS-5 total scores at TP1 (n=132, r=-0.185, p=0.018), indicating that greater severity of post-traumatic symptoms shortly after exposure was associated with decreased individual tendency to take risky choices in the SDRC game (Fig. 4a). This behavioral tendency towards safe behavior was particularly associated with more severe avoidance (CAPS-5 cluster C) (r=-0.244, p=0.003, p_FDR_=0.012) and intrusive symptoms (CAPS-5 cluster B) (r=-0.212, p=0.016, p_FDR_=0.032), but not with other symptom clusters (Fig. 4a). Contrary to our hypothesis, non-significant correlations were found between risky choice index at TP1 and CAPS-5 total scores at TP2 (n=115, r=-0.039, p=0.341) and TP3 (n=112, r=-0.073, p=0.226).

**Figure 4.**
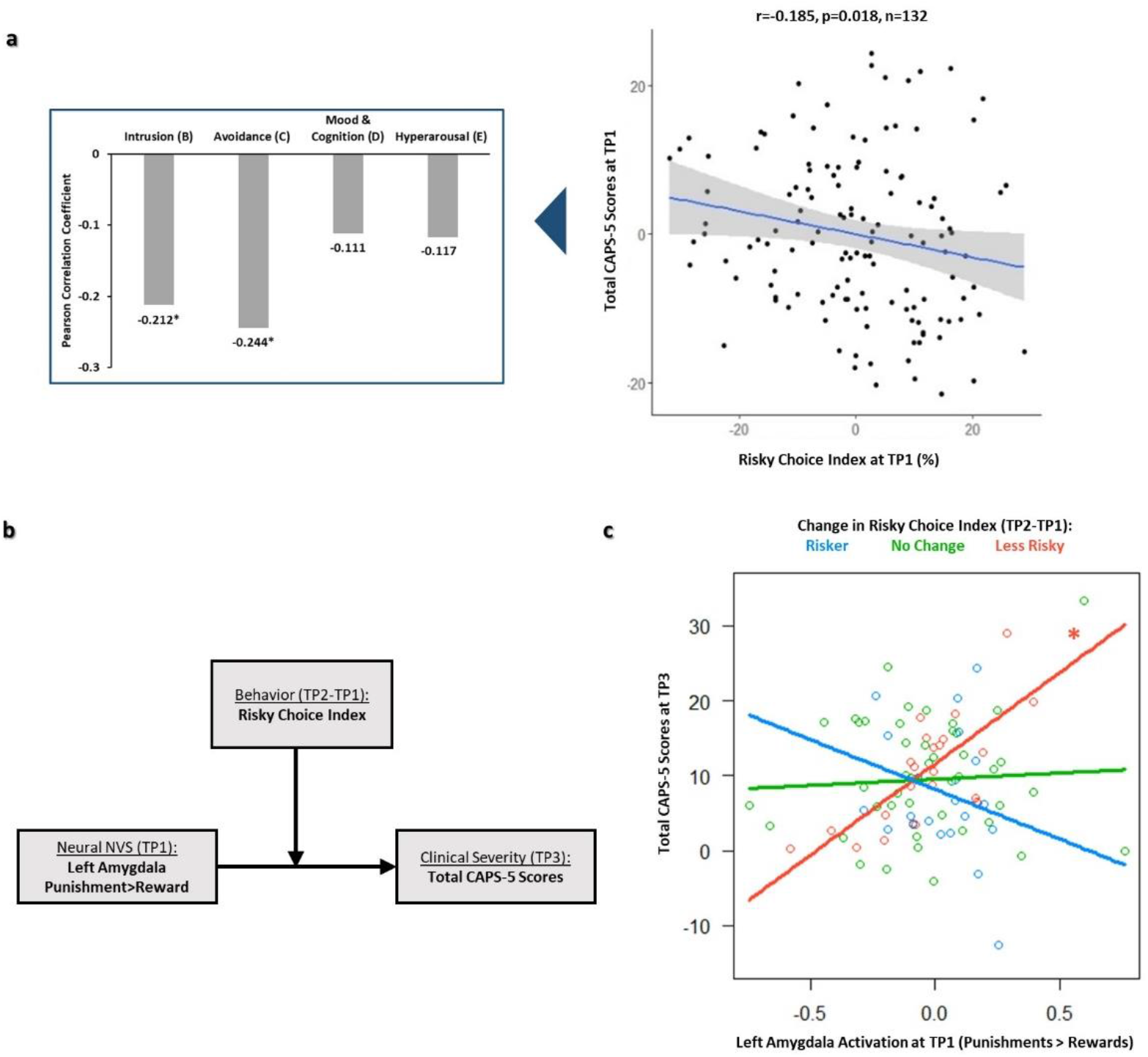
Behavioral Indicators for the Complimentary Role of PVS and NVS in PTSD Symptom Development. **a**. On the right, partial regression plot depicting the relation between individuals’ risky choice index at TP1 (%, x-axis) and their total CAPS-5 scores (y-axis) at TP1, while controlling for age, gender and trauma type (covariates). On the left, a bar plot presenting the correlations between risky choice index and all four PTSD symptom clusters at TP1 according to CAPS-5 (B, C, D, E). Pearson correlation coefficients (r) are presented above each bar. *p-FDR<0.05.**b**. Integrative moderation model of neural NVS, behavior and clinical symptoms: behavioral change in risky choice index (from TP1 to TP2) significantly moderated the relation between valence specific left amygdala activation at TP1 and PTSD symptoms at TP3, beyond age, gender, trauma type and initial symptom severity. **c**. Conditional effects of TP1 left amygdala’s activation to punishments vs. rewards on TP3 CAPS-5 total scores at different values of n=90 individuals’ change in risky choice index (TP2-TP1) (blue = riskier behavior, Mean-1SD; green= no change in risky behavior, Mean; red = less risky behavior, Mean+1SD). All variables were centered prior to the analysis. Change in risky choice index is presented as a categorical variable with 3 levels for illustration purposes, even though it was used as continuous variable in the analyses. *p >.05

In light of the null finding of the relationship between risky choice index at TP1 and PTSD symptoms at TP2 or TP3, we further explored the possibility that behavioral change in risky choice in the SRDC game (TP2-TP1 or TP3-TP1) moderates the relation between early NVS or PVS activity (at TP1) and post-traumatic stress symptom severity 14-months following trauma exposure (TP3). Moderation effects were tested with PROCESS macro for SPSS^71,72^ using hierarchical multiple regression analysis with centered variables and a centered interaction term. Specifically, eight moderation models were tested, including four different neural activations (left and right amygdala and VS at TP1) and two different behavioral measures (change in risky choice index from TP1 to TP2, and from TP1 to TP3). Of these eight moderation models, only one was statistically significant after correcting for multiple comparisons (q_-FDR_<0.05; Fig. 4b). This regression model consisted of left amygdala activation to punishments (vs. rewards) at TP1, change in risky choice index from TP1 to TP2, the interaction between them and four covariates (age, gender, trauma type, and initial symptom severity), together accounting for a significant amount of variance in symptom severity at TP3 (n=90; R^2^=0.400, F_7,82_=7.805, p<0.001). A significant interaction (moderation) effect was detected between amygdala activation at TP1 and change in risk-taking behavior from TP1 to TP2, in predicting CAPS-5 total scores at TP3 (b=-1.074, t_82_=-3.311, p=0.001, p_FDR_=0.011) (see supplementary table 2). This significant interaction was further pursued by testing the conditional effects of left amygdala activation at different values of behavioral change in risky choice index among participants (Mean-1SD, Mean, Mean+1SD) (see Fig. 4c). Interestingly, for individuals that demonstrated a shift towards less risky choices over time (negative change in TP2-TP1 risky choice index, blue line), amygdala activation to punishments at TP1 was associated with more symptoms at TP3 (t=3.754, p=0.003). However, for those with no major change (green line) or with a shift towards riskier behavior from TP1 to TP2 (red line), the relation between amygdala activation at TP1 and CAPS-5 total scores at TP3 was not significant (p>0.05; Fig. 4c).

## Discussion

The longitudinal design of this fMRI study, along with the use of a naturalistic gambling task in a large cohort of recent trauma survivors, enabled a multi-parametric assessment of the relationships between neurobehavioral components of valence systems and PTSD symptom development during the first year following trauma. Negative valence system (NVS) functionality was indicated by amygdala response to punishments vs. rewards, while positive valence system (PVS) functionality was indicated by ventral striatum (VS) response to rewards vs. punishments. Results revealed a differential contribution for these systems in the early manifestation of post-traumatic psychopathology within one month after trauma exposure (TP1) (Fig. 2), as well as in predicting PTSD severity (beyond initial symptoms) over the first year following traumatic exposure (TP3) (Fig 3). Risky choice behavior during the SRDC game revealed the complementary role of both valence systems in PTSD manifestations (Fig. 4). Of note, the neural model of specific brain responses to reward and punishment is a simplification of the human positive and negative valence systems, involving multiple brain areas and networks and the interactions between them^1,73,74^. Future studies may shed additional light on these processes by examining neural responses using a network perspective or data-driven whole-brain approaches.

### NVS (but not PVS) neural functionality is associated with early PTSD severity

The large cohort of symptomatic recent trauma survivors (n=132) recruited from the ER, provided the opportunity to examine the contribution of PVS and NVS neural functioning to PTSD symptom severity as early as one-month after trauma (TP1). Consistent with the vast literature on increased amygdala response to negative stimuli in PTSD^20,27–29,75–77^, hyperactive amygdala to punishments (vs. rewards) during the SRDC game one month after trauma was associated with increased post-traumatic stress symptom severity at that time. Similarly, increased amygdala-lOFC functional connectivity in response to punishments was also linked to greater PTSD severity. The OFC has a known role in the modulation of the amygdala^78,79^ during volitional suppression of negative emotion^80^ and in the presence of threatening stimuli^81^, and its connectivity with the amygdala is involved in processing negative outcomes that signal a need for a behavioral change^82,83^. Along this line, disturbed fronto-amygdalar connectivity was previously observed in PTSD patients^84,85^, but also in patients of other affective psychopathologies^86–90^. Although the current study design cannot disentangle causes from consequences of traumatic stress, the causal role of the NVS functionality was implicated in previous prospective studies, showing for example that hyperactive amygdala in soldiers prior to combat exposure was associated with more PTSD symptoms following exposure^25,91^.

### Contribution of early neural functionality of NVS and PVS to long-term PTSD symptom development

Using a longitudinal design, enabled to demonstrate that NVS and PVS neural functionality shortly after trauma predicted PTSD severity about one year following it. Specifically, excessive amygdala response to punishment and diminished VS response to reward at TP1 were associated with more severe symptoms at TP3 (beyond initial symptom severity) (Fig 3). These opposite relations of the early NVS and PVS with PTSD severity at TP3, allude to a similar finding by Admon et al. (2013)^25^ in a-priori healthy soldiers. This past study showed that increased PTSD symptoms after-exposure to stressful military experience corresponded to increased amygdala response to risk, both pre-and post-exposure, and to decreased VS response to reward, only post-exposure. Likewise, here, the amygdala’s response to punishment at TP1 was related to PTSD symptoms both at TP1 and TP3, whereas VS response to reward at TP1 was only predictive of subsequent symptoms at TP3. Both studies, in line with a putative casual model of PTSD development^91^, suggest that while heightened NVS neural activity represents an early risk factor for this disorder, diminished PVS neural activity may contribute to more long-term consequences following the trauma.

The current study further demonstrated that decreased VS-vmPFC connectivity at TP1 was associated with more severe symptoms at TP3. The vmPFC like the VS is a prominent nodes of the reward circuit, involved in value computations and decision-making processes^92,93^. Human neuroimaging studies have repeatedly demonstrated coincident activation and functional connectivity between these regions during reward processing^94,95^. Animal studies further showing that the vmPFC modulates VS activity^96–98^, and that vmPFC damage was associated with decreased VS volume and diminished response to reward^99^, supporting a causal role of the vmPFC for reward processing via the VS. This corresponds to the results obtained from an explainable machine learning method employing all the used measures of valence systems’ functionality. This analysis revealed that PVS connectivity and activity one month after trauma was more important to the prediction of PTSD development a year later more than NVS activity at this point (Fig. 3f). Together, these findings point to the early role of VS functionality in PTSD severity later on, corresponding to theoretical accounts regarding the importance of PVS in stress recovery by broadening attention and building cognitive and social resources^100,101^.

Our results further showed that PVS and NVS reactivity predicted specific clusters of PTSD symptoms, possibly pointing to distinct underlying mental processes. To this end, hyperactivation within the NVS at TP1 was associated mostly with symptom clusters of hyperarousal and intrusion a year later (CAPS-5 clusters B and E; Fig. 3). These two symptom clusters are well-known as hallmarks of magnified threat processing in PTSD, previously linked to hyperactive NVS in general and amygdala in specific^20–3,102,103^. On the contrary, PVS hypoactivation at TP1 was particularly associated with avoidance symptoms a year later (CAPS-5 cluster C, Fig. 3). Recent animal studies suggested that dopaminergic neurons in the VS regulate approach-avoidance behavior under goal-conflict situations^104,105^. Comparably in humans, VS activity measured by fMRI discriminated between personality tendencies for approach or avoidance under naturalistic high goal-conflict situations^106^. To note, we did not find a correlation between decreased PVS and negative alternations in mood and cognition symptoms (CAPS-5 cluster D), which includes symptoms of anhedonia. In line with findings linking avoidance with deficient reward processing^106–108^, our result suggest that trauma-related avoidance symptoms might be related not only to heightened negative, but also diminished positive valence processing.

Interestingly, amygdala and VS activations at TP1 did not significantly predict PTSD symptom severity at TP2. This null-result might be explained by the dynamic clinical manifestations during the first year following trauma exposure, displaying an overall progressive reduction in PTSD symptom severity, but large inter-individual variability^109–112^. An intermediary point of 6-months post-trauma might be too early to capture the tangible chronic PTSD subtype, whereas 14-months may portray a more stable representation of the chronic disorder, as it was shown to predict over 90% of the expected recovery from PTSD^113,114^. A similar trend of null-results at TP2 was observed in a previous analysis of this data set examining neuroanatomical risk factors for PTSD^115^.

### The potential complementary role of NVS and PVS in post-traumatic psychopathology

We operationalized the risky choice behavior during the SRDC game as an indication of combined involvement of PVS and NVS in the development of PTSD severity. Behavioral results showed that decreased risk-taking during the game at one month after trauma was associated with more severe post-traumatic stress symptoms at the same time-point. This behavioral tendency might reflect increased threat avoidance, potentially due to heightened NVS activity, among individuals with elevated PTSD symptoms in the aftermath of trauma. The rules of the SRDC game encourage a certain amount of risk-taking (e.g., choice of non-matching chip) in order to gain rewards and win (e.g., dispose all domino chips before the time runs out). Therefore, decreased risky behavior at TP1, which was associated with more severe PTSD, stands for reduced likelihood to achieve rewards in light of the risk of receiving punishments. This can be interpreted as over-emphasized NVS and reduced engagement of the PVS in decision-making processes. Such imbalance between the two valence systems is in line with previous work in chronic PTSD patients reporting increased behavioral aversion to risky monetary gains and ambiguous monetary losses^24,26^. This further corresponds to the idea that trauma exposure might alter the homeostatic balance in motivation behavior towards less approach and more avoidance, which in turn might lead to the development of chronic PTSD^15^. The risky choice index in the SRDC game was associated with both intrusion and avoidance symptom clusters of the CAPS-5 (Fig. 4a), the most correlated clusters with the NVS and PVS neural functionality, respectively (Fig. 3 a&b). This combined indication supports our a-priori assertion that reduced risk-taking in the SRDC could represent enhanced NVS along with decreased PVS functionality.

In contrary to our expectations, we did not find a direct association between early risk choice behavior and long-term PTSD severity. Instead, an exploratory moderation analysis showed that a decrease in risky behavior over time (from TP1 to TP2) moderated the relationship between early hyperactive NVS (at TP1) and PTSD symptom development (at TP3) (Fig. 4 b&c). It remains to be determined what drives such temporal dynamics in risk-taking behavior following traumatic stress. Speculatively, a possible mechanism involves cognitive flexibility, the ability to change and adapt one’s behavior in response to changing environments and stimuli^116,117^. Such flexibility may facilitate goal-directed behavior (e.g., approach or avoidance) that promotes the survival and wellbeing of the organism even in the face of danger^118–120^. Previous work has highlighted the importance of early cognitive flexibility compared to other neurocognitive domains, as it was shown to be the most significant cognitive predictor of PTSD symptoms at 14-months post-trauma^121,122^. Furthermore, early neurocognitive intervention improved cognitive flexibility which in turn reduced PTSD symptoms, suggesting its role as a modifiable target preceding and underlying the development of post-traumatic psychopathology^121,123^. Hence, enhancement of cognitive flexibility may enable more risky approach behavior toward prospective rewards, even in the presence of potential threat/punishment, possibly by affecting the interplay between the PVS and NVS. Future research may investigate cognitive flexibility as a potential moderator of the complementary role of the valence systems in development of stress-related psychopathology.

### Conclusion

Using a longitudinal design and a large cohort of recent trauma survivors, the current neuroimaging study was able to provide novel insights on the differential roles of positive and negative valence systems in the development of post-traumatic stress psychopathology. Overall, our findings suggest that the activation and connectivity of NVS and PVS as early as one-month after trauma, could point to specific PTSD symptoms a year later, thus guiding more precise and personalized early intervention. While PTSD research to date has mostly focused on the NVS (e.g., fear, threat), our findings suggest an important (and possibly leading) role for the PVS functionality, in the risk of developing severe symptoms in the first year after trauma. This work also revealed connections between the valence systems functionality and severity in specific symptom clusters, supporting the idea that distinct processes may underlie different clinical phenotypes of PTSD. As the neurobehavioral mechanisms of the human response to positive and negative valence are intrinsically linked, novel therapeutic strategies for PTSD should benefit from addressing symptoms while considering both valence fronts^124^.

## Acknowledgments

This work was supported by award number R01-MH-103287 from the National Institute of Mental Health (NIMH) given to AS (PI), IL and TH (co-Investigators, subcontractors), and had undergone critical review by the NIMH Adult Psychopathology and Disorders of Aging study section. This work was also supported by funding to the Human Brain Project from the European Union Seventh Framework Program (FP7/2007-2013) under grant agreement no. 604102 (HBP). Sagol School of Neuroscience at Tel-Aviv University, Sagol Brain Institute at Tel-Aviv Sourasky Medical Center, and the Human Brain Project (HBP) from the European Union Seventh Framework Program supported the authors’ fellowships. The authors would like to thank the research team at Tel-Aviv Sourasky Medical Center -including Nili Green, Mor Halevi, Sheli Luvton, Yael Shavit, Olga Nevenchannaya, Iris Rashap, Efrat Routledge and Ophir Leshets -for their major contribution in carrying out this research, including subjects’ recruitment and screening, and performing clinical and neural assessments. We also want to extend our gratitude to all the participants of this study, who completed all the assessments at three different time-points after experiencing a traumatic event.

## Ethics

The study was approved by the ethics committee in the local Medical Center (Reference number 0207/14). All participants gave written informed consent in accordance with the Declaration of Helsinki, and received financial remuneration at the end of each time-point (TP1, TP2 and TP3). The study was registered under ClinicalTrials.gov database (registration ID#: NCT03756545; https://clinicaltrials.gov/ct2/show/NCT03756545).

## References

1. Tye KM. Neural Circuit Motifs in Valence Processing. Neuron. 2018;100(2):436–452. doi:10.1016/j.neuron.2018.10.001

2. Insel T, Cuthbert B, Garvey M, et al. Research Domain Criteria (RDoC): Toward a new classification framework for research on mental disorders. Am J Psychiatry. 2010;167(7):748–751. doi:10.1176/appi.ajp.2010.09091379

3. Cuthbert BN, Insel TR. Toward the future of psychiatric diagnosis: The seven pillars of RDoC. BMC Med. 2013;11(1):126. doi:10.1186/1741-7015-11-126

4. Reynolds SM, Berridge KC. Emotional environments retune the valence of appetitive versus fearful functions in nucleus accumbens. Nat Neurosci. 2008;11(4):423–425. doi:10.1038/nn2061

5. Shabel SJ, Wang C, Monk B, Aronson S, Malinow R. Stress transforms lateral habenula reward responses into punishment signals. Proc Natl Acad Sci U S A. 2019;116(25):12488–12493. doi:10.1073/pnas.1903334116

6. Symmonds M, Bossaerts P, Dolan RJ. A behavioral and neural evaluation of prospective decision-making under risk. J Neurosci. 2010;30(43):14380–14389. doi:10.1523/JNEUROSCI.1459-10.2010

7. Levy I, Schiller D. Neural Computations of Threat. Trends Cogn Sci. 2021;25(2):151–171. doi:10.1016/j.tics.2020.11.007

8. Shields GS, Sazma MA, Yonelinas AP. The effects of acute stress on core executive functions: A meta-analysis and comparison with cortisol. Neurosci Biobehav Rev. 2016;68:651–668. doi:10.1016/j.neubiorev.2016.06.038

9. Dreisbach G, Goschke T. How Positive Affect Modulates Cognitive Control: Reduced Perseveration at the Cost of Increased Distractibility. J Exp Psychol Learn Mem Cogn. 2004;30(2):343–353. doi:10.1037/0278-7393.30.2.343

10. Goschke T. Voluntary action and cognitive control from a cognitive neuroscience perspective. In: Voluntary Action: Brains, Minds, and Sociality.; 2003:49–85.

11. Hommel B. Between Persistence and Flexibility. Adv Motiv Sci. 2015:33–67. doi:10.1016/bs.adms.2015.04.003

12. Speer ME, Mauricio R. Reminiscing about positive memories buffers acute stress responses. Nat Hum Behav. 2017;1(5). doi:10.1038/s41562-017-0093

13. Tugade MM, Fredrickson BL. Resilient Individuals Use Positive Emotions to Bounce Back From Negative Emotional Experiences. J Pers Soc Psychol. 2004;86(2):320–333. doi:10.1037/0022-3514.86.2.320

14. McNaughton N, Corr PJ. The neuropsychology of fear and anxiety: A foundation for Reinforcement Sensitivity Theory. In: The Reinforcement Sensitivity Theory of Personality.; 2008:44–94. doi:10.1017/CBO9780511819384.003

15. Stein MB, Paulus MP. Imbalance of Approach and Avoidance: The Yin and Yang of Anxiety Disorders. Biol Psychiatry. 2009;66(12):1072–1074. doi:10.1016/j.biopsych.2009.09.023

16. Weaver SS, Kroska EB, Ross MC, et al. Sacrificing reward to avoid threat: Characterizing PTSD in the context of a trauma-related approach-avoidance conflict task. J Abnorm Psychol. 2020;129(5):457–468. doi:10.1037/abn0000528

17. Aupperle RL, Paulus MP. Neural systems underlying approach and avoidance in anxiety disorders. Dialogues Clin Neurosci. 2010;12(4):517–531. doi:10.31887/dcns.2010.12.4/raupperle

18. Hayes JP, Hayes SM, Mikedis AM, et al. Quantitative meta-analysis of neural activity in posttraumatic stress disorder. Biol Mood Anxiety Disord. 2012;2(1):9. doi:10.1186/2045-5380-2-9

19. Hayes JP, VanElzakker MB, Shin LM. Emotion and Cognition Interactions in PTSD: A Review of Neurocognitive and Neuroimaging Studies. Front Integr Neurosci. 2012;6:89. doi:10.3389/fnint.2012.00089

20. Ronzoni G, del Arco A, Mora F, Segovia G. Enhanced noradrenergic activity in the amygdala contributes to hyperarousal in an animal model of PTSD. Psychoneuroendocrinology. 2016;70:1–9. doi:10.1016/j.psyneuen.2016.04.018

21. Bruce SE, Buchholz KR, Brown WJ, Yan L, Durbin A, Sheline YI. Altered emotional interference processing in the amygdala and insula in women with Post-Traumatic Stress Disorder. NeuroImage Clin. 2013;2(1):43–49. doi:10.1016/j.nicl.2012.11.003

22. Hayes JP, LaBar KS, McCarthy G, et al. Reduced hippocampal and amygdala activity predicts memory distortions for trauma reminders in combat-related PTSD. J Psychiatr Res. 2011;45(5):660–669. doi:10.1016/j.jpsychires.2010.10.007

23. Forster GL, Simons RM, Baugh LA. Revisiting the Role of the Amygdala in Posttraumatic Stress Disorder. In: The Amygdala - Where Emotions Shape Perception, Learning and Memories.; 2017. doi:10.5772/67585

24. Jia R, Ruderman L, Gordon C, et al. A Shift from Value- to Saliency-Neural Encoding of Subjective Value in Combat Veterans with PTSD during Decision Making under Uncertainty. bioRxiv. 2020:2020.04.14.041467. doi:10.1101/2020.04.14.041467

25. Admon R, Lubin G, Rosenblatt JD, et al. Imbalanced neural responsivity to risk and reward indicates stress vulnerability in humans. Cereb Cortex. 2013;23(1):28–35. doi:10.1093/cercor/bhr369

26. Ruderman L, Ehrlich DB, Roy A, Pietrzak RH, Harpaz-Rotem I, Levy I. Posttraumatic Stress Symptoms and Aversion To Ambiguous Losses in Combat Veterans. Depress Anxiety. 2016;33(7):606–613. doi:10.1002/da.22494

27. Zuj D V., Palmer MA, Lommen MJJ, Felmingham KL. The centrality of fear extinction in linking risk factors to PTSD: A narrative review. Neurosci Biobehav Rev. 2016;69:15–35. doi:10.1016/j.neubiorev.2016.07.014

28. LeDoux JE. Emotion circuits in the brain. Annu Rev Neurosci. 2000;23:155–184. doi:10.1146/annurev.neuro.23.1.155

29. Graham BM, Milad MR. The study of fear extinction: Implications for anxiety disorders. Am J Psychiatry. 2011;168(12):1255–1265. doi:10.1176/appi.ajp.2011.11040557

30. Aghajani M, Veer IM, van Hoof M-J, Rombouts SARB, van der Wee NJ, Vermeiren RRJM. Abnormal functional architecture of amygdala-centered networks in adolescent posttraumatic stress disorder. Hum Brain Mapp. 2016;37(3):1120–1135. doi:10.1002/hbm.23093

31. Zhu X, Suarez-Jimenez B, Lazarov A, et al. Exposure-based therapy changes amygdala and hippocampus resting-state functional connectivity in patients with posttraumatic stress disorder. Depress Anxiety. 2018;35(10):974–984. doi:10.1002/da.22816

32. Nawijn L, van Zuiden M, Frijling JL, Koch SBJ, Veltman DJ, Olff M. Reward functioning in PTSD: A systematic review exploring the mechanisms underlying anhedonia. Neurosci Biobehav Rev. 2015;51:189–204. doi:10.1016/j.neubiorev.2015.01.019

33. Seidemann R, Duek O, Jia R, Levy I, Harpaz-Rotem I. The Reward System and Post-Traumatic Stress Disorder: Does Trauma Affect the Way We Interact With Positive Stimuli? Chronic Stress. 2021;5:247054702199600. doi:10.1177/2470547021996006

34. Ikemoto S. Brain reward circuitry beyond the mesolimbic dopamine system: A neurobiological theory. Neurosci Biobehav Rev. 2010;35(2):129–150. doi:10.1016/j.neubiorev.2010.02.001

35. Nikolova YS, Bogdan R, Brigidi BD, Hariri AR. Ventral striatum reactivity to reward and recent life stress interact to predict positive affect. Biol Psychiatry. 2012;72(2):157–163. doi:10.1016/j.biopsych.2012.03.014

36. Pizzagalli DA, Holmes AJ, Dillon DG, et al. Reduced caudate and nucleus accumbens response to rewards in unmedicated individuals with major depressive disorder. Am J Psychiatry. 2009;166(6):702–710. doi:10.1176/appi.ajp.2008.08081201

37. Stoy M, Schlagenhauf F, Sterzer P, et al. Hyporeactivity of ventral striatum towards incentive stimuli in unmedicated depressed patients normalizes after treatment with escitalopram. J Psychopharmacol. 2012;26(5):677–688. doi:10.1177/0269881111416686

38. Elman I, Lowen S, Frederick BB, Chi W, Becerra L, Pitman RK. Functional Neuroimaging of Reward Circuitry Responsivity to Monetary Gains and Losses in Posttraumatic Stress Disorder. Biol Psychiatry. 2009;66(12):1083–1090. doi:10.1016/j.biopsych.2009.06.006

39. Sailer U, Robinson S, Fischmeister FPS, et al. Altered reward processing in the nucleus accumbens and mesial prefrontal cortex of patients with posttraumatic stress disorder. Neuropsychologia. 2008;46(11):2836–2844. doi:10.1016/j.neuropsychologia.2008.05.022

40. Felmingham KL, Falconer EM, Williams L, et al. Reduced amygdala and ventral striatal activity to happy faces in PTSD is associated with emotional numbing. PLoS One. 2014;9(9). doi:10.1371/journal.pone.0103653

41. Mehta ND, Stevens JS, Li Z, et al. Inflammation, reward circuitry and symptoms of anhedonia and PTSD in trauma-exposed women. Soc Cogn Affect Neurosci. 2020. doi:10.1093/scan/nsz100

42. Zhu X, Helpman L, Papini S, et al. Altered resting state functional connectivity of fear and reward circuitry in comorbid PTSD and major depression. Depress Anxiety. 2017. doi:10.1002/da.22594

43. American Psychiatric Association, Association AP. Diagnostic And Statistical Manual of Mental Disorders?: DSM-5. American Psychiatric Association; 2013. https://ci.nii.ac.jp/naid/10010095609/.

44. Rab SL, Admon R. Parsing inter- and intra-individual variability in key nervous system mechanisms of stress responsivity and across functional domains. Neurosci Biobehav Rev. 2020. doi:10.1016/j.neubiorev.2020.09.007

45. Galatzer-Levy IR, Bryant RA. 636,120 Ways to Have Posttraumatic Stress Disorder. Perspect Psychol Sci. 2013;8(6):651–662. doi:10.1177/1745691613504115

46. Young G, Lareau C, Pierre B. One Quintillion Ways to Have PTSD Comorbidity: Recommendations for the Disordered DSM-5. Psychol Inj Law. 2014;7(1):61–74. doi:10.1007/s12207-014-9186-y

47. Pitman RK, Rasmusson AM, Koenen KC, et al. Biological studies of post-traumatic stress disorder. Nat Rev Neurosci. 2012;13(11):769–787. doi:10.1038/nrn3339

48. Zoladz P, Diamond D. Current status on behavioral and biological markers of PTSD. Neurosci Biobehav Rev. 2013.

49. Ben-Zion Z, Fine NB, Keynan NJ, et al. Neurobehavioral Moderators of Post-Traumatic Stress Disorder Trajectories: Prospective fMRI Study of Recent Trauma Survivors. Eur J Psychotraumatol. 2019;10(1). doi:10.1080/20008198.2019.1683941

50. Shalev AY, Gevonden M, Ratanatharathorn A, et al. Estimating the risk of PTSD in recent trauma survivors: results of the International Consortium to Predict PTSD (ICPP). World Psychiatry. 2019;18(1):77–87. doi:10.1002/wps.20608

51. Shalev AY, Ankri Y, Israeli-Shalev Y, Peleg T, Adessky R, Freedman S. Prevention of posttraumatic stress disorder by early treatment: Results from the Jerusalem trauma outreach and prevention study. Arch Gen Psychiatry. 2012;69(2):166–176. doi:10.1001/archgenpsychiatry.2011.127

52. Kahn I, Yeshurun Y, Rotshtein P, Fried I, Ben-Bashat D, Hendler T. The role of the amygdala in signaling prospective outcome of choice. Neuron. 2002;33(6):983–994.

53. Assaf M, Kahn I, Pearlson GD, et al. Brain activity dissociates mentalization from motivation during an interpersonal competitive game. Brain Imaging Behav. 2009;3(1):24–37. doi:10.1007/s11682-008-9047-y

54. Gonen T, Admon R, Podlipsky I, Hendler T. From animal model to human brain networking: Dynamic causal modeling of motivational systems. J Neurosci. 2012;32(21):7218–7224. doi:10.1523/JNEUROSCI.6188-11.2012

55. Thaler A, Gonen T, Mirelman A, et al. Altered reward-related neural responses in non-manifesting carriers of the Parkinson disease related LRRK2 mutation. Brain Imaging Behav. 2019;13(4):1009–1020. doi:10.1007/s11682-018-9920-2

56. Admon R, Lubin G, Stern O, et al. Human vulnerability to stress depends on amygdala’s predisposition and hippocampal plasticity. Proc Natl Acad Sci. 2009;106(33):14120–14125. doi:10.1073/pnas.0903183106

57. Blanchard EB, Jones-Alexander J, Buckley TC, Forneris CA. Psychometric properties of the PTSD checklist (PCL). Behav Res Ther. 1996;34(8):669–673. doi:10.1016/0005-7967(96)00033-2

58. Weathers FW, Bovin MJ, Lee DJ, et al. The Clinician-Administered PTSD Scale for DSM–5 (CAPS-5): Development and initial psychometric evaluation in military veterans. Psychol Assess. 2018;30(3):383–395. doi:10.1037/pas0000486

59. Hyatt CJ, Assaf M, Muska CE, et al. Reward-related dorsal striatal activity differences between former and current cocaine dependent individuals during an interactive competitive game. PLoS One. 2012;7(5):1–15. doi:10.1371/journal.pone.0034917

60. Esteban O, Markiewicz CJ, Blair RW, et al. fMRIPrep: a robust preprocessing pipeline for functional MRI. Nat Methods. 2019;16(1):111–116. doi:10.1038/s41592-018-0235-4

61. Gorgolewski K, Burns CD, Madison C, et al. Nipype: A flexible, lightweight and extensible neuroimaging data processing framework in Python. Front Neuroinform. 2011;5. doi:10.3389/fninf.2011.00013

62. Fan L, Li H, Zhuo J, et al. The Human Brainnetome Atlas: A New Brain Atlas Based on Connectional Architecture. Cereb Cortex. 2016;26(8):3508–3526. doi:10.1093/cercor/bhw157

63. Pauli WM, Nili AN, Michael Tyszka J. Data Descriptor: A high-resolution probabilistic in vivo atlas of human subcortical brain nuclei. Sci Data. 2018;5. doi:10.1038/sdata.2018.63

64. Brett M, Anton JL, Valabregue R, Poline JB. Region of interest analysis using an SPM toolbox. Neuroimage. 2002.

65. McLaren DG, Ries ML, Xu G, Johnson SC. A generalized form of context-dependent psychophysiological interactions (gPPI): A comparison to standard approaches. Neuroimage. 2012. doi:10.1016/j.neuroimage.2012.03.068

66. Whitfield-Gabrieli S, Nieto-Castanon A. Conn: A Functional Connectivity Toolbox for Correlated and Anticorrelated Brain Networks. Brain Connect. 2012;2(3):125–141. doi:10.1089/brain.2012.0073

67. IBM Corp. IBM SPSS Statistics for Windows, Version 26.0. 2019. 2019.

68. R Core Team (2020). R: A language and environment for statistical computing. R A Lang Environ Stat Comput R Found Stat Comput Vienna, Austria. 2020.

69. Benjamini Y, Hochberg Y. Controlling the False Discovery Rate: A Practical and Powerful Approach to Multiple Testing. J R Stat Soc Ser B. 1995;57(1):289–300. doi:10.1111/j.2517-6161.1995.tb02031.x

70. Lundberg SM, Lee SI. A unified approach to interpreting model predictions. In: Advances in Neural Information Processing Systems. Vol 2017-Decem.; 2017:4766–4775.

71. Preacher KJ, Hayes AF. Asymptotic and resampling strategies for assessing and comparing indirect effects in multiple mediator models. Behav Res Methods. 2008;40(3):879–891. doi:10.3758/BRM.40.3.879

72. Hayes AF. Introduction to Mediation, Moderation and Conditional Process Analysis. Vol 53.; 2013. doi:10.5539/ass.v11n9p207

73. Berridge KC. Affective valence in the brain: modules or modes? Nat Rev Neurosci. 2019;20(4):225–234. doi:10.1038/s41583-019-0122-8

74. Baxter MG, Murray EA. The amygdala and reward. Nat Rev Neurosci. 2002;3(7):563–573. doi:10.1038/nrn875

75. Liberzon I, Sripada CS. The functional neuroanatomy of PTSD: a critical review. Prog Brain Res. 2007;167:151–169. doi:10.1016/S0079-6123(07)67011-3

76. Shin LM, Liberzon I. The Neurocircuitry of Fear, Stress and Anxiety Disorders. Neuropsychopharmacology. 2010;35(1):169–191. doi:10.1038/npp.2009.83

77. Stevens JS, Kim YJ, Galatzer-Levy IR, et al. Amygdala Reactivity and Anterior Cingulate Habituation Predict Posttraumatic Stress Disorder Symptom Maintenance After Acute Civilian Trauma. Biol Psychiatry. 2017;81(12):1023–1029. doi:10.1016/j.biopsych.2016.11.015

78. Amiel Rosenkranz J, Grace AA. Cellular mechanisms of infralimbic and prelimbic prefrontal cortical inhibition and dopaminergic modulation of basolateral amygdala neurons in vivo. J Neurosci. 2002;22(1):324–337. doi:10.1523/jneurosci.22-01-00324.2002

79. Quirk GJ, Likhtik E, Pelletier JG, Paré D. Stimulation of medial prefrontal cortex decreases the responsiveness of central amygdala output neurons. J Neurosci. 2003;23(25):8800–8807. doi:10.1523/jneurosci.23-25-08800.2003

80. Phan KL, Fitzgerald DA, Nathan PJ, Moore GJ, Uhde TW, Tancer ME. Neural substrates for voluntary suppression of negative affect: A functional magnetic resonance imaging study. Biol Psychiatry. 2005;57(3):210–219. doi:10.1016/j.biopsych.2004.10.030

81. Garcia R, Vouimba RM, Baudry M, Thompson RF. The amygdala modulates prefrontal cortex activity relative to conditioned fear. Nature. 1999;402(6759):294–296. doi:10.1038/46286

82. Elliott R, Dolan RJ, Frith CD. Dissociable functions in the medial and lateral orbitofrontal cortex: Evidence from human neuroimaging studies. Cereb Cortex. 2000;10(3):308–317. doi:10.1093/cercor/10.3.308

83. O’Doherty J, Kringelbach ML, Rolls ET, Hornak J, Andrews C. Abstract reward and punishment representations in the human orbitofrontal cortex. Nat Neurosci. 2001;4(1):95–102. doi:10.1038/82959

84. Shin LM, Wright CI, Cannistraro PA, et al. A functional magnetic resonance imaging study of amygdala and medial prefrontal cortex responses to overtly presented fearful faces in posttraumatic stress disorder. Arch Gen Psychiatry. 2005;62(3):273–281. doi:10.1001/archpsyc.62.3.273

85. Williams LM, Kemp AH, Felmingham K, et al. Trauma modulates amygdala and medial prefrontal responses to consciously attended fear. Neuroimage. 2006;29(2):347–357. doi:10.1016/j.neuroimage.2005.03.047

86. Hahn A, Stein P, Windischberger C, et al. Reduced resting-state functional connectivity between amygdala and orbitofrontal cortex in social anxiety disorder. Neuroimage. 2011;56(3):881–889. doi:10.1016/j.neuroimage.2011.02.064

87. Monk CS, Telzer EH, Mogg K, et al. Amygdala and ventrolateral prefrontal cortex activation to masked angry faces in children and adolescents with generalized anxiety disorder. Arch Gen Psychiatry. 2008;65(5):568–576. doi:10.1001/archpsyc.65.5.568

88. Frodl T, Bokde ALW, Scheuerecker J, et al. Functional Connectivity Bias of the Orbitofrontal Cortex in Drug-Free Patients with Major Depression. Biol Psychiatry. 2010;67(2):161–167. doi:10.1016/j.biopsych.2009.08.022

89. Drevets WC, Price JL, Furey ML. Brain structural and functional abnormalities in mood disorders: Implications for neurocircuitry models of depression. Brain Struct Funct. 2008;213(1-2):93–118. doi:10.1007/s00429-008-0189-x

90. Versace A, Thompson WK, Zhou D, et al. Abnormal Left and Right Amygdala-Orbitofrontal Cortical Functional Connectivity to Emotional Faces: State Versus Trait Vulnerability Markers of Depression in Bipolar Disorder. Biol Psychiatry. 2010;67(5):422–431. doi:10.1016/j.biopsych.2009.11.025

91. Admon R, Milad MR, Hendler T. A causal model of post-traumatic stress disorder: disentangling predisposed from acquired neural abnormalities. Trends Cogn Sci. 2013;17(7):337–347. doi:10.1016/J.TICS.2013.05.005

92. Pauli WM, O’Reilly RC, Yarkoni T, Wager TD. Regional specialization within the human striatum for diverse psychological functions. Proc Natl Acad Sci U S A. 2016;113(7):1907–1912. doi:10.1073/pnas.1507610113

93. Clithero JA, Rangel A. Informatic parcellation of the network involved in the computation of subjective value. Soc Cogn Affect Neurosci. 2013;9(9):1289–1302. doi:10.1093/scan/nst106

94. Cauda F, Cavanna AE, D’agata F, Sacco K, Duca S, Geminiani GC. Functional connectivity and coactivation of the nucleus accumbens: A combined functional connectivity and structure-based meta-analysis. J Cogn Neurosci. 2011;23(10):2864–2877. doi:10.1162/jocn.2011.21624

95. Choi EY, Thomas Yeo BT, Buckner RL. The organization of the human striatum estimated by intrinsic functional connectivity. J Neurophysiol. 2012;108(8):2242–2263. doi:10.1152/jn.00270.2012

96. Voorn P, Vanderschuren LJMJ, Groenewegen HJ, Robbins TW, Pennartz CMA. Putting a spin on the dorsal-ventral divide of the striatum. Trends Neurosci. 2004;27(8):468–474. doi:10.1016/j.tins.2004.06.006

97. Gabbott PLA, Warner TA, Jays PRL, Salway P, Busby SJ. Prefrontal cortex in the rat: Projections to subcortical autonomic, motor, and limbic centers. J Comp Neurol. 2005;492(2):145–177. doi:10.1002/cne.20738

98. Ghazizadeh A, Ambroggi F, Odean N, Fields HL. Prefrontal cortex mediates extinction of responding by two distinct neural mechanisms in accumbens shell. J Neurosci. 2012;32(2):726–737. doi:10.1523/JNEUROSCI.3891-11.2012

99. Pujara MS, Philippi CL, Motzkin JC, Baskaya MK, Koenigs M. Ventromedial prefrontal cortex damage is associated with decreased ventral striatum volume and response to reward. J Neurosci. 2016;36(18):5047–5054. doi:10.1523/JNEUROSCI.4236-15.2016

100. Fredrickson BL. What good are positive emotions? Rev Gen Psychol. 1998. doi:10.1037/1089-2680.2.3.300

101. Fredrickson BL, Tugade MM, Waugh CE, Larkin GR. What good are positive emotions in crisis? J Pers Soc Psychol. 2003.

102. Rabellino D, Densmore M, Frewen PA, Théberge J, Lanius RA. The innate alarm circuit in post-traumatic stress disorder: Conscious and subconscious processing of fear- and trauma-related cues. Psychiatry Res - Neuroimaging. 2016;248:142–150. doi:10.1016/j.pscychresns.2015.12.005

103. Lanius RA, Rabellino D, Boyd JE, Harricharan S, Frewen PA, McKinnon MC. The innate alarm system in PTSD: conscious and subconscious processing of threat. Curr Opin Psychol. 2017;14:109–115. doi:10.1016/j.copsyc.2016.11.006

104. Nguyen D, Fugariu V, Erb S, Ito R. Dissociable roles of the nucleus accumbens D1 and D2 receptors in regulating cue-elicited approach-avoidance conflict decision-making. Psychopharmacology (Berl). 2018;235(8):2233–2244. doi:10.1007/s00213-018-4919-3

105. Nguyen D, Alushaj E, Erb S, Ito R. Dissociative effects of dorsomedial striatum D1 and D2 receptor antagonism in the regulation of anxiety and learned approach-avoidance conflict decision-making. Neuropharmacology. 2019;146:222–230. doi:10.1016/j.neuropharm.2018.11.040

106. Gonen T, Soreq E, Eldar E, Ben-Simon E, Raz G, Hendler T. Human mesostriatal response tracks motivational tendencies under naturalistic goal conflict. Soc Cogn Affect Neurosci. 2016;11(6):961–972. doi:10.1093/scan/nsw014

107. Romanczuk-Seiferth N, Koehler S, Dreesen C, Wüstenberg T, Heinz A. Pathological gambling and alcohol dependence: Neural disturbances in reward and loss avoidance processing. Addict Biol. 2015;20(3):557–569. doi:10.1111/adb.12144

108. Simon JJ, Walther S, Fiebach CJ, et al. Neural reward processing is modulated by approach- and avoidance-related personality traits. Neuroimage. 2010;49(2):1868- doi:10.1016/j.neuroimage.2009.09.016

109. Hepp U, Moergeli H, Buchi S, et al. Post-traumatic stress disorder in serious accidental injury: 3-Year follow-up study. Br J Psychiatry. 2008;192(5):376–383. doi:10.1192/bjp.bp.106.030569

110. Yehuda R, McFarlane AC, Shalev AY. Predicting the development of posttraumatic stress disorder from the acute response to a traumatic event. Biol Psychiatry. 1998;44(12):1305–1313. doi:10.1016/S0006-3223(98)00276-5

111. Koren D, Arnon I, Klein E. Long term course of chronic posttraumatic stress disorder in traffic accident victims: A three-year prospective follow-up study. Behav Res Ther. 2001;39(12):1449–1458. doi:10.1016/S0005-7967(01)00025-0

112. Perkonigg A, Pfister H, Stein MB, et al. Longitudinal course of posttraumatic stress disorder and posttraumatic stress disorder symptoms in a community sample of adolescents and young adults. Am J Psychiatry. 2005;162(7):1320–1327. doi:10.1176/appi.ajp.162.7.1320

113. Shalev AY, Freedman S. PTSD Following Terrorist Attacks: A Prospective Evaluation. Am J Psychiatry. 2005;162(6):1188–1191. doi:10.1176/appi.ajp.162.6.1188

114. Kessler RC, Sonnega A, Bromet E, Hughes M, Nelson CB. Posttraumatic Stress Disorder in the National Comorbidity Survey. Arch Gen Psychiatry. 1995;52(12):1048–1060. doi:10.1001/archpsyc.1995.03950240066012

115. Ben-Zion Z, Artzi M, Niry D, et al. Neuroanatomical Risk Factors for Post Traumatic Stress Disorder (PTSD) in Recent Trauma Survivors. Biol Psychiatry Cogn Neurosci Neuroimaging. 2019:721134. doi:10.1101/721134

116. Cañas JJ, Quesada JF, Antolí A, Fajardo I. Cognitive flexibility and adaptability to environmental changes in dynamic complex problem-solving tasks. Ergonomics. 2003;46(5):482–501. doi:10.1080/0014013031000061640

117. Cañ JJ, Antoli A, Fajardo I, Salmerón L. Cognitive inflexibility and the development and use of strategies for solving complex dynamic problems: Effects of different types of training. Theor Issues Ergon Sci. 2005;6(1):95–108. doi:10.1080/14639220512331311599

118. Berridge KC, Kringelbach ML. Neuroscience of affect: Brain mechanisms of pleasure and displeasure. Curr Opin Neurobiol. 2013;23(3):294–303. doi:10.1016/j.conb.2013.01.017

119. Berridge KC, Kringelbach ML. Pleasure Systems in the Brain. Neuron. 2015;86(3):646–664. doi:10.1016/j.neuron.2015.02.018

120. Lang PJ, Bradley MM. Emotion and the motivational brain. Biol Psychol. 2010;84(3):437–450. doi:10.1016/j.biopsycho.2009.10.007

121. Ben-Zion Z, Fine NB, Keynan NJ, et al. Cognitive Flexibility Predicts PTSD Symptoms: Observational and Interventional Studies. Front Psychiatry. 2018;9:477. doi:10.3389/fpsyt.2018.00477

122. Ben-Zion Z, Zeevi Y, Keynan NJ, et al. Multi-domain potential biomarkers for post-traumatic stress disorder (PTSD) severity in recent trauma survivors. Transl Psychiatry. 2020. doi:10.1038/s41398-020-00898-z

123. Joseph J, Moring J, Bira L. Cognitive Flexibility as a Key Factor in the Conceptualization and Treatment of PTSD. Curr Psychiatry Rev. 2015. doi:10.2174/1573400511666150629104921

124. Dutcher JM, Creswell JD. The role of brain reward pathways in stress resilience and health. Neurosci Biobehav Rev. 2018;95:559–567. doi:10.1016/j.neubiorev.2018.10.014

