## Supplementary Materials for "Differential Roles of Positive and Negative Valence Systems in the Development of Post-Traumatic Stress Psychopathology"

Supplementary Methods

Supplementary Results

Supplementary Table 1. Whole-brain Activations to Rewards vs. Punishments.

Supplementary Table 2. Change in risky choice index moderates the relationship between amygdala's response to punishments at TP1 and PTSD symptom severity at TP3.

Supplementary Figure 1. Whole-brain Activations to Rewards vs. Punishments.

**Supplementary Methods**

**fMRI Paradigm.** Before the MRI scan, outside the scanner, participants were instructed regarding the aim and rules of the fMRI domino paradigm. They also performed a short training session in order to verify that they understood the task's instructions. During the scan, individuals played for 14 minutes using a 4-buttons remote controller for moving left, right and selecting a specific domino chip. From our experience, most participants found this task very engaging and seemed to believe the idea that they were actually plating against the experimenter.

**fMRI Data Analysis.** Raw DICOM data images were converted to NIFTI format and organized to conform to the 'Brain Imaging Data Structure' specifications (BIDS)[2]. Preprocessing was conducted using FMRIPREP version 1.5.8[3], a Nipype based tool[4].
*(i) Anatomical Preprocessing.* Within the FMRIPREP framework, each of T1-weighted (T1w) image was corrected for intensity non-uniformity (INU) using `N4BiasFieldCorrection' v2.1.0, distributed with `AntsApplyTransforms` (ANTs version 2.2.0). The T1w reference was then skull-stripped with a Nipype implementation of the `antsBrainExtraction.sh` workflow (from ANTs), using OASIS30-ANTs as a target template. Brain tissue segmentation of cerebrospinal fluid (CSF), white matter (WM) and gray matter (GM) was performed on the brain-extracted T1w using `FAST` (FSL version 5.0.9). A T1w-reference map was computed after registration of the INU-corrected T1w image using `mri_robust_template` (FreeSurfer version 6.0.1). Volume-based spatial normalization to one standard space (MNI152NLin2009cAsym) was performed through nonlinear registration with `antsRegistration` tool of ANTs version 2.2.0, using brain-extracted versions of both T1w reference and the T1w template. The ICBM 152 nonlinear Asymmetrical template version 2009 was selected for spatial normalization.
*(ii) Functional Preprocessing.* First, a reference volume and its skull-stripped version were generated using a custom methodology of FMRIPREP, and the susceptibility distortion correction (SDC) was omitted. The BOLD reference was then co-registered to the T1w reference using `MCFLIRT` (FSL version 5.0.9) with the boundary-based registration cost-function. Co-registration was configured with nine degrees of freedom to account for distortions remaining in the BOLD reference. Head-motion parameters with respect to the BOLD reference (transformation matrices, and six corresponding rotation and translation parameters) were estimated before any spatio-temporal filtering. BOLD runs were slice-time corrected using `3dTshift` from AFNI version 16.2.07, and their time-series were resampled onto their original, native space by applying the transforms to correct for head-motion. Several confounding time-series were calculated based on framewise displacement (FD), DVARS and three region-wise global signals (extracted within the CSF, the WM, and the whole-brain masks). Additionally, a set of physiological regressors were

extracted to allow for component-based noise correction (CompCor). Principal components were estimated after high-pass filtering of the pre-processed BOLD time-series (using a discrete cosine filter with 128s cut-off) for the two CompCor variants: temporal (tCompCor) and anatomical (aCompCor). Six tCompCor components were then calculated including only the top 5% variable voxels within that subcortical mask. For aCompCor, six components were calculated within the intersection of the subcortical mask and the union of CSF and WM masks calculated in T1w space, after their projection to the native space of each functional run. For each CompCor decomposition, the k components with the largest singular values were retained, sufficient to explain 50% of variance across the nuisance mask. The remaining components were dropped from consideration. The head-motion estimates calculated in the correction step were also placed within the corresponding confounds file. The confound time series derived from head motion estimates and global signals were expanded with the inclusion of temporal derivatives and quadratic terms. Frames that exceeded a threshold of 0.5mm FD or 1.5 standardized DVARS were annotated as motion outliers. All re-samplings was performed with a single interpolation step by composing all the pertinent transformations. Gridded (volumetric) re-samplings were performed using ANTs, configured with Lanczos interpolation to minimize the smoothing effects of other kernels, while non-gridded (surface) re-samplings were performed using `mri_vol2surf` (FreeSurfer). Many internal operations of FMRIPREP use `Nilearn`, principally within the BOLD-processing workflow (for details, refer to https://fmriprep.readthedocs.io/en/stable/workflows.html). Finally, spatial smoothing of the data was performed using SPM12 (full-width at half-maximum: 6mm).

*Whole-Brain and ROI Analysis.* When participant had no actual choice, when only matching or only non-matching chips remained to choose from, the events were classified as false rounds ("false"). When participant did not respond within the defined time to choose a chip (during 10 seconds starting with the "go" signal), the events were also classified as false rounds ("false"). When participant had to move and choose a chip (during both "ready" and "go" intervals), the events were classified as movements. Implicit baseline was defined as both decision-making intervals ("choose") and false rounds ("false"). For the whole-brain analysis, Significance level was set at p=0.05, with whole brain with family-wise error (FWE) correction. For results, please refer to supplementary results.

*Functional Connectivity Analysis.* After preprocessing steps using FMRIPREP functional MRI data was further preprocessed for functional connectivity analysis using the CONN toolbox (version 18.b)[5]. The default denoising pipeline was performed, combining both linear regression of potential confounding effects in the BOLD signal, and temporal band-pass filtering. Denoising pipeline included two steps. The first step was linear regression, in which factors that were identified as potential confounding effects to the estimated BOLD signal were estimated and removed separately for each voxel and for each subject and functional run/session. This was done using Ordinary Least Squares (OLS) regression to project each BOLD signal time series to the sub-space orthogonal to all potential confounding effects. Potential confounding effects used in CONN's default denoising pipeline implement an anatomical component-based noise correction procedure (aCompCor), and include noise components from cerebral white matter and cerebrospinal areas[6], estimated subject-motion parameters[7], identified outlier scans or scrubbing[8], constant and first-order linear session effects, and constant task effects[5]. The second step was temporal band-pass filtering, in which temporal frequencies below 0.008 Hz were removed from the BOLD signal in order to focus on slow-frequency fluctuations, while minimizing the influence of physiological, head-motion and other noise sources. Filtering was implemented using a discrete cosine transform windowing operation to minimize border effects, and performed after regression to avoid any frequency mismatch in the nuisance regression procedure[9]. Then, functional connectivity analysis was performed using generalized psychophysiological interaction (gPPI) as implemented in CONN toolbox[10]. Unlike the standard PPI analysis that includes contrast information when forming a psychological regressor, the gPPI approach convolves the BOLD signal with the canonical hemodynamic response function for each condition before making the contrast, forming a separate psychological regressor for each condition. This approach has been known to improve the fit of the regression model for event-related fMRI data[10], hence fits the analysis of the domino task described here. First, we extracted an average BOLD time-course across selected voxels for each ROI and used it as a physiological regressor. Second, we computed how strongly the time-course of one ROI is correlated with the PPI regressor of another. This was done using a separate multiple regression model for each target ROI time-series. Third, results were converted to z-scores using the Fisher's z-transformation before calculating a group-level averaged FC. Again, the examined contrast here (as in all other analyses throughout this work) was both rewards vs. both punishments.

### Supplementary Results

#### Neurobehavioral indicators of PVS and NVS for PTSD severity shortly after trauma

At the neural level, whole-brain analysis for the rewards vs. punishments contrast yielded several clusters at $p<0.05$ FWE corrected, including in the VS (see Supplementary Figure 1 and Supplementary Table 1). The opposite contrast of punishments vs. rewards yielded no significant clusters.

#### Neurobehavioral predictors of PTSD development in the first year after trauma

To note, the neural activations were not bilateral; while left VS activation to reward at TP1 was marginally significantly correlated with PTSD symptom severity at TP3 *(n=111, r=-0.159, p=0.051)*, right amygdala activation to punishment at TP1 was not associated with PTSD symptom severity at TP3 *(n=108; r=0.082, p=0.204)*.

To note, significant correlations were also found between TP3 symptom severity and TP1 connectivity patterns of right VS-left vmPFC *(r=-0.216, p=0.027)* and left VS-right vmPFC *(r=-0.208, p=0.034)*, but these did not survive correction for multiple comparisons *(for both: $q_{\text{-FDR}}>0.05$)* (not shown).

### Supplementary Tables

**Supplementary Table 1. Whole-brain Activations to Rewards vs. Punishments.** Brain regions revealed by whole-brain regression analysis in the contrast of rewards vs. punishments (thresholded at T=5.958, p<.0005 FWE whole-brain corrected, k>50 voxels). L/R=Peak in Left/Right hemisphere; k=cluster size; T=T-score; MNI coordinates =x,y,z voxel coordinates in MNI space of the peak voxel.

| Brain region | L/R | k | T | MNI coordinates | | |
|---|---|---|---|---|---|---|
| | | | | x | y | z |
| Striatum (Ventral & Dorsal) | R | 163 | 11.86 | 16 | 10 | -7 |
| | R | | 11.24 | 11 | 16 | -1 |
| | R | | 7.14 | 16 | 19 | 9 |
| | L | 133 | 11.58 | -13 | 10 | -4 |
| Inferior Parietal Lobule | R | 174 | 8.92 | 37 | -37 | 45 |
| | | | 7.74 | 49 | -31 | 45 |
| | L | 57 | 7.35 | -46 | -49 | 39 |
| Inferior Frontal Gyrus | R | 54 | 7.33 | 52 | 4 | 24 |
| | | | 6.78 | 52 | 13 | 36 |

**Supplementary Table 2. Change in risky choice index moderates the relationship between amygdala's response to punishments at TP1 and PTSD symptom severity at TP3.** Regression model of n=90 participants' CAPS-5 total scores at TP3 predicted from: left amygdala's average activation to punishments (vs. rewards) at TP1; change in risky choice index from TP1 to TP2; the interaction between these two variables; participants' age, gender, trauma type and initial symptom severity (CAPS-5 total scores at TP1). For each variable, standardized coefficient beta, t-value, and two-tailed significance level of t –value are specified in the table. *p <.05

| **Predictor** | **β** | **t** | ***p*** |
|---|---|---|---|
| Amygdala's Activation to Punishments (TP1) | 2.55 | 0.73 | .497 |
| Change in Risky Choice Index (TP2 – TP1) | -0.10 | -1.64 | .105 |
| **Interaction*** | -1.07 | -3.31 | .001 |
| Age (covariate) | 0.06 | 0.76 | .447 |
| Gender (covariate) | -0.07 | -0.04 | .969 |
| Trauma Type (covariate) | -0.18 | -0.11 | .913 |
| **Total CAPS-5 at TP1*** (covariate) | 0.46 | 6.16 | .000 |

### Supplementary Figures

**Supplementary Figure 1. Whole-brain Activations to Rewards vs. Punishments.** Statistical parametric T-maps (sagittal, coronal and axial) obtained from all TP1 participants (n=132) showing brain regions with increased activation in response to both rewards vs. both punishments ($p<0.05$, FWE whole-brain corrected). The color bar at the bottom represents positive T-value, ranging from 2.893 (black) to 11 (white). No negative T-values (de-activations) were observed for this significance level. See supplementary Table 1 for a list of full activations.

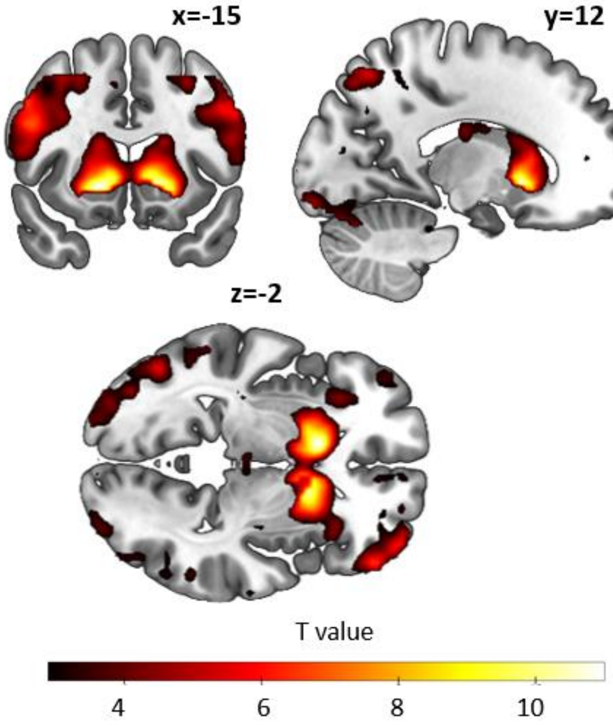

**References**


1. Ben-Zion Z, Fine NB, Keynan NJ, et al. Neurobehavioral moderators of post-traumatic stress disorder (PTSD) trajectories: study protocol of a prospective MRI study of recent trauma survivors. *Eur J Psychotraumatol*. 2019;10(1). doi:10.1080/20008198.2019.1683941

2. Gorgolewski KJ, Auer T, Calhoun VD, et al. The brain imaging data structure, a format for organizing and describing outputs of neuroimaging experiments. *Sci Data*. 2016;3. doi:10.1038/sdata.2016.44

3. Esteban O, Markiewicz CJ, Blair RW, et al. fMRIPrep: a robust preprocessing pipeline for functional MRI. *Nat Methods*. 2019;16(1):111-116. doi:10.1038/s41592-018-0235-4

4. Gorgolewski K, Burns CD, Madison C, et al. Nipype: A flexible, lightweight and extensible neuroimaging data processing framework in Python. *Front Neuroinform*. 2011;5. doi:10.3389/fninf.2011.00013

5. Whitfield-Gabrieli S, Nieto-Castanon A. Conn: A Functional Connectivity Toolbox for Correlated and Anticorrelated Brain Networks. *Brain Connect*. 2012;2(3):125-141. doi:10.1089/brain.2012.0073

6. Behzadi Y, Restom K, Liau J, Liu TT. A component based noise correction method (CompCor) for BOLD and perfusion based fMRI. *Neuroimage*. 2007;37(1):90-101. doi:10.1016/j.neuroimage.2007.04.042

7. Friston KJ, Williams S, Howard R, Frackowiak RSJ, Turner R. Movement-related effects in fMRI time-series. *Magn Reson Med*. 1996;35(3):346-355. doi:10.1002/mrm.1910350312

8. Power JD, Mitra A, Laumann TO, Snyder AZ, Schlaggar BL, Petersen SE. Methods to detect, characterize, and remove motion artifact in resting state fMRI. *Neuroimage*. 2014;84:320-341. doi:10.1016/j.neuroimage.2013.08.048

9. Hallquist MN, Hwang K, Luna B. The nuisance of nuisance regression: Spectral misspecification in a common approach to resting-state fMRI preprocessing reintroduces noise and obscures functional connectivity. *Neuroimage*. 2013;82:208-225. doi:10.1016/j.neuroimage.2013.05.116

10. McLaren DG, Ries ML, Xu G, Johnson SC. A generalized form of context-dependent psychophysiological interactions (gPPI): A comparison to standard approaches. *Neuroimage*. 2012. doi:10.1016/j.neuroimage.2012.03.068
